# Citrus genomic resources unravel putative genetic determinants of Huanglongbing, a pathogen-triggered immune disease

**DOI:** 10.1101/2022.10.24.513527

**Authors:** Yuxia Gao, Jin Xu, Zhilong Li, Yunzeng Zhang, Nadia Riera, Zhiwei Xiong, Zhigang Ouyang, Xinjun Liu, Zhanjun Lu, Danelle Seymour, Balian Zhong, Nian Wang

## Abstract

Citrus is one of the most important tree crops worldwide and citrus production is threatened by Huanglongbing (HLB), a devasting citrus disease caused by *Candidatus* Liberibacter. HLB is a pathogen-triggered immune disease, in which the pathogen initiates systemic and chronic immune responses including the excessive production of reactive oxidative species, which subsequently lead to cell death of phloem tissues and HLB disease symptoms. Here, we identified putative genetic determinants of HLB pathogenicity by integrating citrus genomic resources to characterize the pan-genome of accessions that differ in their response to HLB. Genome-wide association mapping and analysis of allele-specific expression between susceptible, tolerant, and resistant accessions further refined candidates underlying the response to HLB. To enable these analyses we first developed a phased diploid assembly of *Citrus sinensis* ‘Newhall’ genome and produced resequencing data for 91 citrus accessions that differ in their response to HLB. These data were combined with previous resequencing data from 356 sequenced accessions for genome-wide association mapping of the HLB response. Genes with HLB pathogenicity were associated with the host immune response, ROS production, and antioxidants. Overall, this study has provided a significant recourse of citrus genomic data and we have identified candidate genes to be further explored to understand the genetic determinants of HLB pathogenicity and to generate HLB resistant/tolerant citrus varieties.

## Introduction

Citrus is one of top three fruit crops worldwide and is an important source of vitamin C in human diet (Wu et al., 2014). However, citrus production faces many challenges including diseases, drought, flood, and freezes. Among them, citrus Huanglongbing (HLB, also known as greening) caused by *Candidatus* Liberibacter spp. presents an unprecedented challenge and has spread to most citrus growing regions such as China, Brazil, and USA (**Bové**, 2006; Pandey et al., 2022). *Ca*. L. asiaticus (CLas) is the most prevalent HLB pathogen. Almost all commercial citrus varieties are susceptible to HLB with few exceptions, such as Sugar Belle Mandarin (*C. reticulata*), Persian lime (*C. latifolia*), and US-897 (*Citrus reticulata* Blanco × *Poncirus trifoliata* L. Raf.) that show tolerance against HLB (Albrecht and Bowman, 2012; Deng et al., 2019; Folimonova et al., 2009; Sivager et al., 2021). In addition, multiple citrus relatives such as *Microcitrus australis, Eremocitrus glauca, Swinglea glutinosa, M. warburgiana*, and *M. papuana*, have shown resistance against HLB (Alves et al., 2020; Cifuentes-Arenas et al., 2019).

HLB has been suggested to be a pathogen-triggered immune disease (Ma et al., 2022). CLas is vectored mainly by Asian citrus psyllid (*Diaphorina citri*) and after transmission begins to colonize the phloem where it initiates a systemic and chronic immune response including reactive oxygen species (ROS) production, subsequent cell death of phloem tissues, and eventual HLB symptom development. Consistent with this model, CLas causes fewer changes in the expression of immunity genes in the tolerant rootstock variety US-897 than the susceptible variety ‘Cleopatra’ mandarin (Albrecht and Bowman, 2012). In addition, CLas stimulates significantly higher levels of ROS production in both the HLB susceptible Mexican lime and the HLB-tolerant Persian lime, with the latter demonstrating higher antioxidants, suggesting that high ROS levels are tolerated in Persian lime (Sivager *et al*., 2021). However, the genetic determinants responsible for resistance, tolerance, and susceptibility of different citrus genotypes against HLB remains unknown.

With HLB being a pathogen-triggered immune disease, it has been proposed that CLas recognition and downstream immune signaling, ROS production, and antioxidants may be responsible for variation in the response to HLB in citrus (resistance, tolerance, and susceptibility) (Ma *et al*., 2022). Antimicrobial peptides have also been associated with resistance to HLB (Huang et al., 2021).

Bacterial pathogens trigger immune response via recognition by either membrane-localized pattern recognition receptors (PRRs) and intracellular nucleotide-binding domain leucine-rich repeat receptors (NLRs) (Ge et al., 2022; Lu and Tsuda, 2021), leading to PAMP-triggered immunity (PTI) and effector-triggered immunity (ETI), respectively. PTI and ETI share multiple downstream immune responses including the influx of Ca^2+^, ROS burst, activation of mitogen-activated protein kinase (MAPK) cascades, defense gene induction, and biosynthesis of defense phytohormones (Ge *et al*., 2022; Lu and Tsuda, 2021). Recent studies showed that both PTI and ETI are needed to mount a robust immune response, as they synergistically enhance each other (Chang et al., 2022; Ngou et al., 2021; Pruitt et al., 2021; Tena, 2021; Tian et al., 2021; Yuan et al., 2021; Zhai et al., 2022). In citrus there are a large number of genes involved in these two signaling pathways. For example, citrus immune signaling cascades include approximately 925 pattern recognition receptors (PRRs), 703 nucleotide-binding, leucine-rich domains (NLRs), 45 calcium-dependent protein kinases (CDPKs or CPKs), 100 mitogen-activated protein kinase (MAPKs), 119 cytoplasmic receptor like kinases (CRLK), as well as 137 pathogenesis-related (PR) genes, 19 ROS genes, and 436 antioxidant enzyme genes. We hypothesized that the genetic determinants of HLB pathogenicity, including candidates in these immune signaling pathways, can be identified through the integration of existing and newly produced citrus genomic resources to facilitate pan-genome analysis, a genome-wide association study (GWAS) for HLB response, and allelic-specific expression analyses of citrus accessions that differ in HLB resistance, tolerance and susceptibility. The genomes of multiple citrus genotypes and relatives have been assembled including *C. sinensis* (Xu et al., 2013), *Atlantia buxifolia, C. medica, C. ichangensis* (Wang et al., 2017), *Fortunella hindsii* (Zhu et al., 2019), *C. clementina* (Wu *et al*., 2014), *C. reticulata* (Wang et al., 2018a), *C. grandis*, and *Poncirus trifoliata* (Peng et al., 2020). However, chromosome-level phased genomes were not available except *C. sinensis* cv. Valencia (Wu et al. 2022), limiting exploration of haplotype-specific differences in gene content. In addition, thousands of accessions of citrus cultivars and relatives are available worldwide, representing a treasure to be mined.

In this study, we developed a haplotype-resolved genome assembly of *C. sinensis* ‘Newhall’, which was used for identification of genes that are specific to HLB resistant accessions. In total, pan-genome analysis was performed using genome assemblies from both susceptible and tolerant/resistant citrus cultivars. In addition, 26 citrus accessions (HLB resistant, tolerant, or susceptible) that have high quality genome sequencing data were mined for small indels linked to HLB response. The phased chromosome-level genome of *C. sinensis* was also used to investigate the difference in HLB susceptibility and tolerance of Valencia sweet orange and Sugar Belle mandarin LB9-9, respectively. We have also sequenced 91 citrus accessions, which, together with 356 previously sequenced citrus accessions, were used for GWAS analysis of genetic determinants responsible for HLB pathogenicity. Overall, this study has developed a significant resource of citrus genomic information and identified candidate genes to be further explored to understand the genetic determinants of HLB pathogenicity and to generate HLB resistant/tolerant citrus varieties.

## Results

### Phased diploid genome assembly of *C. sinensis* ‘Newhall’

We have selected *C. sinensis* ‘Newhall’ for sequencing because of its interesting features including maturation in the winter, whereas most other sweet orange cultivars mature before the summer, a second “twin” fruit opposite its stem (navel), and seedlessness. As a side note, it is worth mentioning that this project was started in 2019, which has suffered many hurdles caused by COVID19 similar as others. In total, 54 gigabases (Gb) Illumina paired-end short read data, 35 Gb PacBio HIFI read data, and 45 Gb Hi-C data were produced (Table S1). The genome of Newhall navel orange was de novo assembled using both HIFI and Illumina reads. The de novo assembly length was 685.27 Mb, including 1,624 contigs with N50 value of 12.5 Mb. Those contigs were further assembled into 1,013 scaffolds with N50 value of 32.8 M (Table S2) using Hi-C scaffolding methods. The final assembly contained 618 Mb in 18 chromosomes, which were assigned into primary or secondary haplotypes. Only 67 Mb sequences were unassigned to a scaffold (Fig. 1, Table S3 and S4). Syntenic blocks between homologous chromosomes indicated that the haplotypes were phased correctly (Fig. S1)

**Figure 1.**
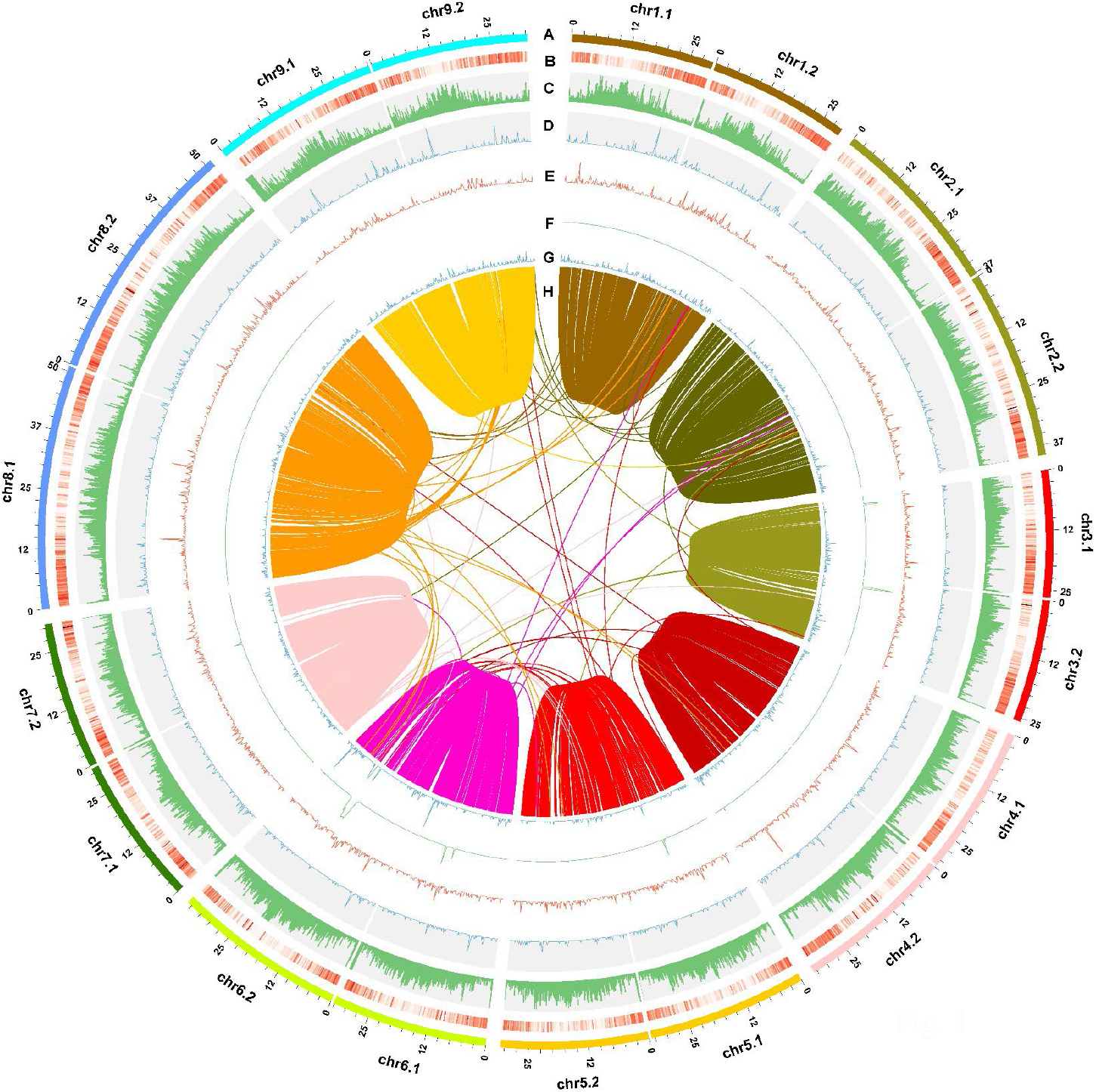
Genomic features of the *Citrus sinensis* Osbeck cv. Newhall. The circle graph depicts the genomic characteristics, including chromosome length (A), gene density (B), density of transposable element (C), density of miRNA (D), density of snRNA (E), density of rRNA (F), density of tRNA (G), and the synteny between homologous chromosomes (H).

The quality of assembly was supported by the *k*-mer analysis, which indicated that the haploid genome size was 350.15 Mb, approximately half of the assembled diploid genome size (Tables S2, and S5). In addition, BUSCO (95.1% of completeness), CEGMA (97.74% of completeness) assessments, and genomic sequencing coverage (99.97%) indicated high integrity of the phased diploid sweet orange genome assembly (Table S6-S8). Synteny blocks were identified and revealed an overall synteny between the homoeologous chromosomes in Newhall navel orange (Fig. 1, Fig. S1-S2). Relatively low similarity values (37.41% to 46.82%) were observed between the homoeologous chromosomes in Newhall navel orange (Table S9).

A total of 356 Mb (52%) of the assembled genome was masked and annotated as repeated sequences, of which 44.42% were long terminal repeat (LTR) retrotransposons, 1.74% were DNA transposable elements, 0.83% were long interspersed nuclear elements (LINE), and 0.06% were short interspersed nuclear elements (SINE) (Fig. 1, Tables S2 and S10). To reduce false positives in gene prediction, we combined de novo, homology, and RNA-seq based approaches (Fig. S3). Consequently, a total of 46,616 gene models were identified (Tables S2 and S11, Fig. S3), among which 45,431 (97.46% of total predicted genes) were protein-coding genes (Table S12). The Newhall genome contained 22,916 genes on one chromosome set and 22,824 genes on the other chromosome set, and 876 genes unable to be assigned to either chromosome set. The annotated genes were classified into 26 COG categories including signal transduction, transcription, post-translational modification, protein turnover, and chaperones, carbohydrate transport and metabolism, translation, ribosomal structure, and biogenesis, intracellular trafficking, secretion, and vesicular transport, and secondary metabolites biosynthesis, transport, and catabolism (Table S13), with more than 42.6% of predicted genes as functionally unknown. In addition, 11,221 non-coding RNA sequences were identified and annotated, including 931 micro RNAs (miRNAs), 1,200 transfer RNAs (tRNAs), 6,765 ribosomal RNA (rRNAs) and 2,325 small nuclear RNA (snRNA) (Fig. 1, Table S14). We found most homologous gene pairs (13,257/15,423) have a *Ka/Ks* ratio <1, indicative of purifying selection (Fig. S4). A total of 894 gene pairs with a *Ka/Ks* ratio >1 may have undergone positive selection, which included genes involved in metabolic and biosynthetic process, and response to stress and stimulus (Fig. S5).

### Pan-genome analyses of genes that are differentially present in HLB tolerant and susceptible genotypes

We explored the level of presence/absence sequence variation across the genus citrus and its relatives to dissect the genetic determinants of HLB pathogenicity. This was done using 9 previously assembled genomes of citrus and relatives and the newly sequenced Newhall genome. The 10 citrus accessions and relatives were classified into HLB-susceptible (6 citrus accessions) and -tolerant groups (4 citrus accessions) (Table S15). Here we have defined HLB-tolerant trees as those showing vigorous growth (such as thick canopy and not dying) in the presence of CLas and HLB symptoms, whereas HLB-susceptible trees refer to those without vigorous growth (such as with thin canopy and dying) in the presence of CLas and HLB symptoms based on the description in the original publications. Pan-genome analyses of the 10 genomes identified 50,442 gene families. The total number of gene sets continued to increase with the addition of each genome and was approaching, but did not reach a plateau at n = 10 (Fig. 2A), suggesting more high quality genomes of citrus and its relatives are required. Specifically,13,301, 4,559, 12,135, and 20,447 gene families were defined as core, soft-core, dispensable, and private genes, respectively (Fig. 2B). The fraction of core genes (core plus soft core) in the citrus pan-genome (35%) was in line with previous studies (35– 87%)(Contreras-Moreira et al., 2017; Gao et al., 2019; Golicz et al., 2016; Gordon et al., 2017; Hurgobin et al., 2018; Li et al., 2014; Montenegro et al., 2017; Sun et al., 2020; Wang et al., 2018b). The dispensable and private gene families accounted for 64.6% of the total pan gene families, enlarging gene resources of citrus reference genome. The core gene families accounted for more than 51.7% genes of each genome (Fig. 2C), suggesting conserved genomic features among citrus and relatives. In addition, wild citrus species, such as Ichang papeda, citron, and kumquat, and primitive citrus *Atlantia*, showed higher proportion of private genes than cultivated citrus accessions (Fig. 2C). More than 47.2% of pan genome gene families were functionally unknown, which was higher than the diploid genome of *C. sinensis* ‘Newhall’ (42.3%) (Fig. S6, Table S13, and Table S16).

**Figure 2.**
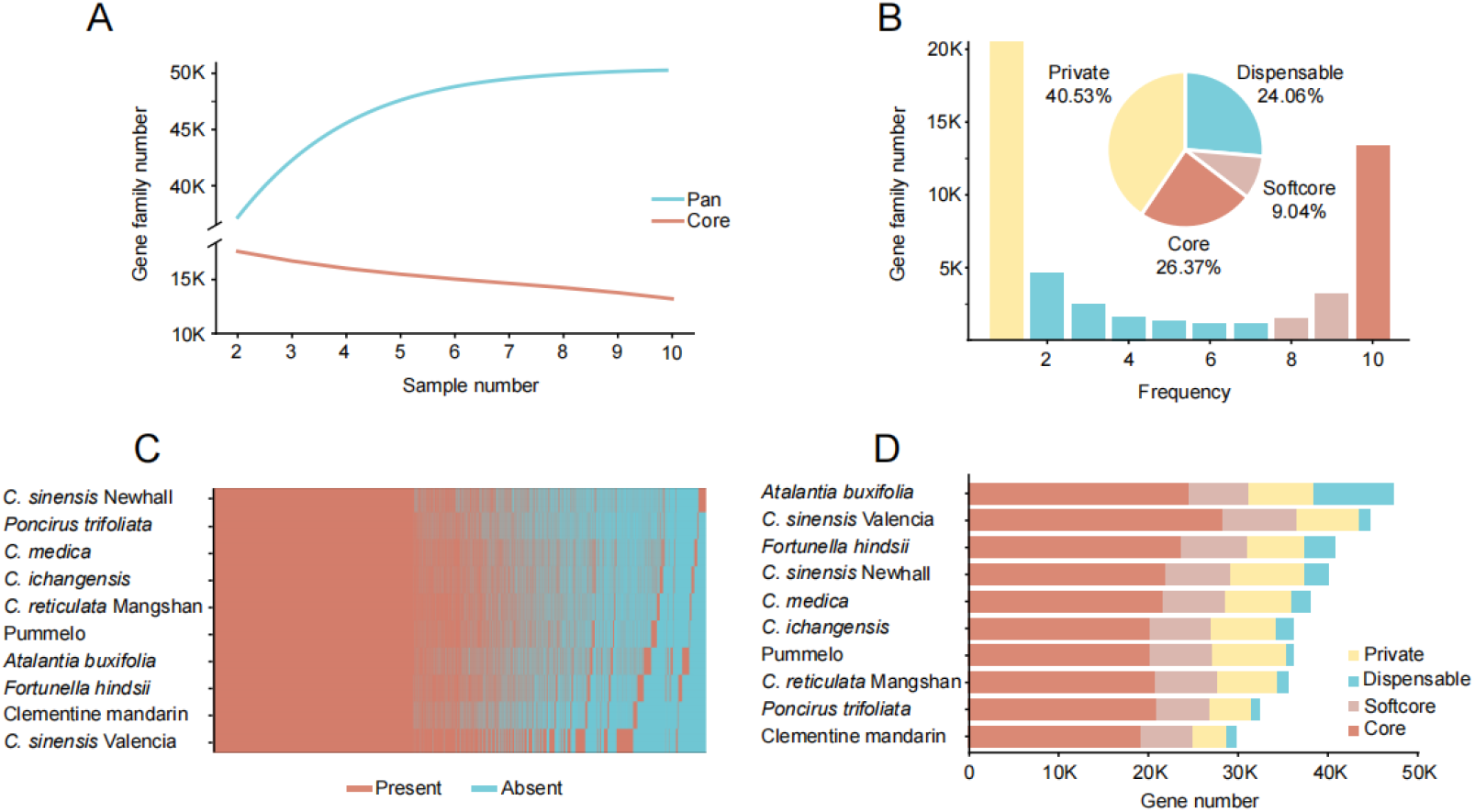
Pan genome of *Citrus* and relatives. a. Rarefaction curve of detected genes in the pan and core genomes. b. The composition of the pan genome, including core, soft core, dispensable and private genes (pie plot). The histogram depicting the number of gene families of pan genome under different presence frequency in 26 accessions of *Citrus* and relatives. c. Presence and absence information of pan gene families in accessions of citrus and citrus relatives. d. Gene number of each composition for each individual genome. HKC: *Atalantia buxifolia*,, CSV: ‘Valencia’ sweet orange, FOR: *Fortunella hindsii*, CSN: ‘Newhall’ sweet orange, XZ: *Citrus medica*, XJC: *Citrus ichangensis*, HWB: pummelo, CMS: *Citrus reticulata* ‘Mangshan’, PON: *Poncirus trifoliata*, and CLM: Clementine mandarin.

Whole-genome alignment between the *C. sinensis* Valencia genome and each of the 9 other genome assemblies was performed. A total of 77,609 InDels (37,355 deletions, 40,254 insertions) were identified in addition to 3 million single nucleotide polymorphisms (SNPs) (Fig. S7, Table S17). We searched genes with presence/absence variation between the HLB tolerant and susceptible groups (Table S15). Since HLB is a pathogen-triggered immune disease (Ma *et al*., 2022), we paid close attention to immunity related genes. The genes families involved in plant immunity, such as CDPK, MAPK, NBS-LRR, NPR, hormone, PLCP, PR, RLKs, ROS, and antioxidant genes, were presented mostly in dispensable and private gene sets (Fig. 3A). Furthermore, HLB-susceptible citrus accessions were enriched for the plant immunity genes that were absent in HLB-tolerant accessions (Fig. 3B, Table S18), suggesting a possible explanation for the stronger systemic and chronic immune responses observed in susceptible citrus accessions compared to those that are tolerant.

**Figure 3.**
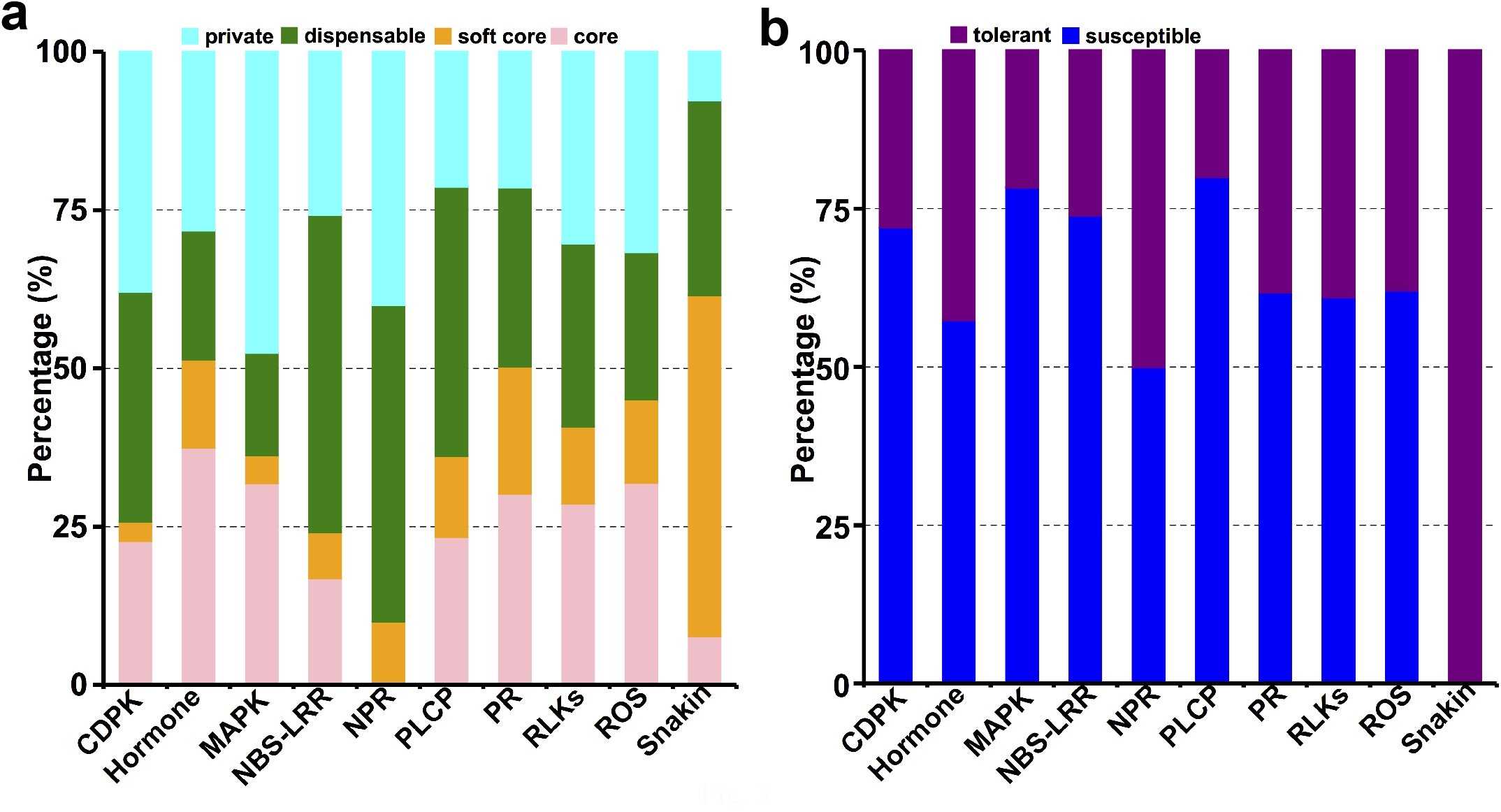
Plant immunity-related genes in citrus pan genome. a. The composition of plant immunity-related genes in citrus pan genome, including core, soft core, dispensable and private genes. b. The HLB-tolerant (including resistant accessions) and -susceptible citrus accessions specific plant immunity-related genes. The plant immunity-related genes here refer to CDPK (Calcium-Dependent Protein Kinases), MAPK (Mitogen-Activated Protein Kinase Cascades), NBS-LRR (Nucleotide Binding Domain and Leucine-rich Repeat), NPR (Nonexpressor of Pathogenesis-related Genes), PLCP (Papain-Like Cysteine Proteases), PR (Pathogenesis-related Protein), RLKs (Receptor-Like Kinases), ROS (Reactive Oxygen Species), snakin, and plant hormone.

We further analyzed group (HLB-tolerant and -susceptible)-specific genes related to immunity (Table S15). Most of immunity related genes were present in all the accessions. Nevertheless, we have identified multiple interesting genes including orange1.1t03332.1 (NBS-LRR), orange1.1t04682.1 (NBS-LRR), orange1.1t05285.1 (PLCP, cysteine protease-like protein), Cs6g22310.1 (lectin), orange1.1t05183.1 (Leucine-rich repeat receptor-like protein kinase), Cs1g05340.1 (LRR-XII), Cs9g13810.1 (RLCK-XII/XIII), and Cs6g09910.5 (MAPKKK, Raf31), which were present in most HLB susceptible accessions (67-83%), but were absent in all HLB resistant accessions. On the contrary, only Cs2g10550.1 (Leucine-rich repeat receptor-like protein kinase), and Cs1g05370.1 (Serine-threonine protein kinase, plant-type) were present in 75% of four HLB tolerant accessions, but absent in the 6 HLB susceptible accessions. In addition, 74 antioxidant biosynthesis and antioxidant enzyme genes were absent in susceptible citrus accessions, but were present in one or more tolerant citrus accessions (Dataset 1).

### Indel analyses of genes involved in plant immunity in 26 HLB-resistant, -tolerant, or - susceptible citrus accessions

The pan-genome analysis of 10 assembled citrus genomes included only accessions susceptible or tolerant to HLB, but none with HLB resistance (refers to citrus plants with no HLB symptoms in the presence of CLas or inhibiting CLas growth). To search for genomic signatures associated with resistance to HLB we expanded our interspecific analysis of genomic variation to a panel of 26 citrus accessions and relatives with members exhibiting all three classes of response to HLB infection. The additional 16 accessions were selected based on their phylogenetic relationships and high quality genome resequencing data (20 to 126 x coverages) (Table S15). In total, we have identified 263177, 26891, and 1369 indels in the resistant, tolerant, and susceptible groups, respectively. Specifically, we identified indels in the coding region of 26 NBS-LRR, 3 receptor-like kinase, and 1 SOD genes in the resistant group, 4 NBS-LRR and 1 SOD genes in the tolerant group, 1 NBS-LRR and 1 receptor-like kinase genes in the susceptible group (Table S19). Consistent with the model that HLB is a pathogen-triggered immune disease (Ma *et al*., 2022), HLB resistant/tolerant citrus accessions contain more indel mutations in plant immunity genes than HLB susceptible accessions that might contribute to the reduced immune responses in tolerant/resistant citrus genotypes compared to the susceptible genotypes (Albrecht and Bowman, 2012; Sivager *et al*., 2021).

### GWAS analysis of citrus genes that are potentially responsible for the HLB resistance/ tolerance or susceptibility

GWAS has been widely used to understand the genetic basis of plant disease resistance and susceptibility. To further explore the genetic basis of HLB resistance, we performed GWAS analysis using a large panel of both HLB susceptible and resistant/tolerant varieties (Fig. 4, Table S20, Dataset 2) using whole genome sequences of 447 citrus accessions and relatives (McMahon et al., 2021). We have sequenced 91 additional citrus genotypes (Tables S21) to complement public sequence data sets of 356 accessions. A total of 7.59 million SNPs for 447 citrus accessions and relatives were generated, which were subsequently filtered by quality and sequence depth, and SNPs with minor allele frequency (MAF) more than 0.01 and individual level missingness less than 10%. A total of 252,357 SNPs across the whole genome were used for HLB GWAS analysis (Fig S8). The principal component analysis based on SNP data suggested 447 citrus accessions and relatives showed population stratification, which was accounted in GWAS analysis (Fig S9). The Quantile-Quantile Plot suggested the robustness of our GWAS analysis (Fig S10). We found HLB-associated citrus genomic SNPs were from a large number of genes across whole genome (Fig. 4a), suggesting that the citrus genetic effect on HLB may be explained by the omnigenic model (a large number of genes). Such a model indicates that complex traits are influenced by core genes with direct effects as well as by a modest number of genes or pathways with small effects (Boyle, Li, and Pritchard 2017). 252 SNPs, including 86 from coding region and 166 from non-coding region were significantly associated with HLB resistance/tolerance and susceptibility (Fig. 4a, Dataset 2, adjusted p-value <1e-5). 37% of genes (32/86) containing SNPs with significant association with HLB resistance/tolerance and susceptibility were involved in plant immunity, ROS, and stress response (Fig. 4b, Dataset 2). However, no group-specific SNPs were identified for HLB resistant/tolerant and susceptible groups even though SNPs from 10 genes showed different patterns between HLB susceptible and tolerant/resistant accessions (Fig. 5, Dataset 2). These 10 genes included *PCS1* (*Phytochelatin Synthase1*), mutation of which impairs callose deposition, bacterial pathogen defense and auxin content (De Benedictis et al., 2018; Morita-Yamamuro et al., 2005); CNGC1 (Cyclic nucleotide-gated ion channel 1) that acts as Ca^2+^-permeable channels involved in Ca^2+^ oscillations and receptor-mediated signaling during plant immunity (Dietrich et al., 2020); two RLKs, *pabAB* (para-aminobenzoate synthetase) which has been shown to scavenge the reactive oxygen species in vitro (Lu et al., 2014); and *RPPL1* (NLB RPP13-like protein 1).

**Figure 4.**
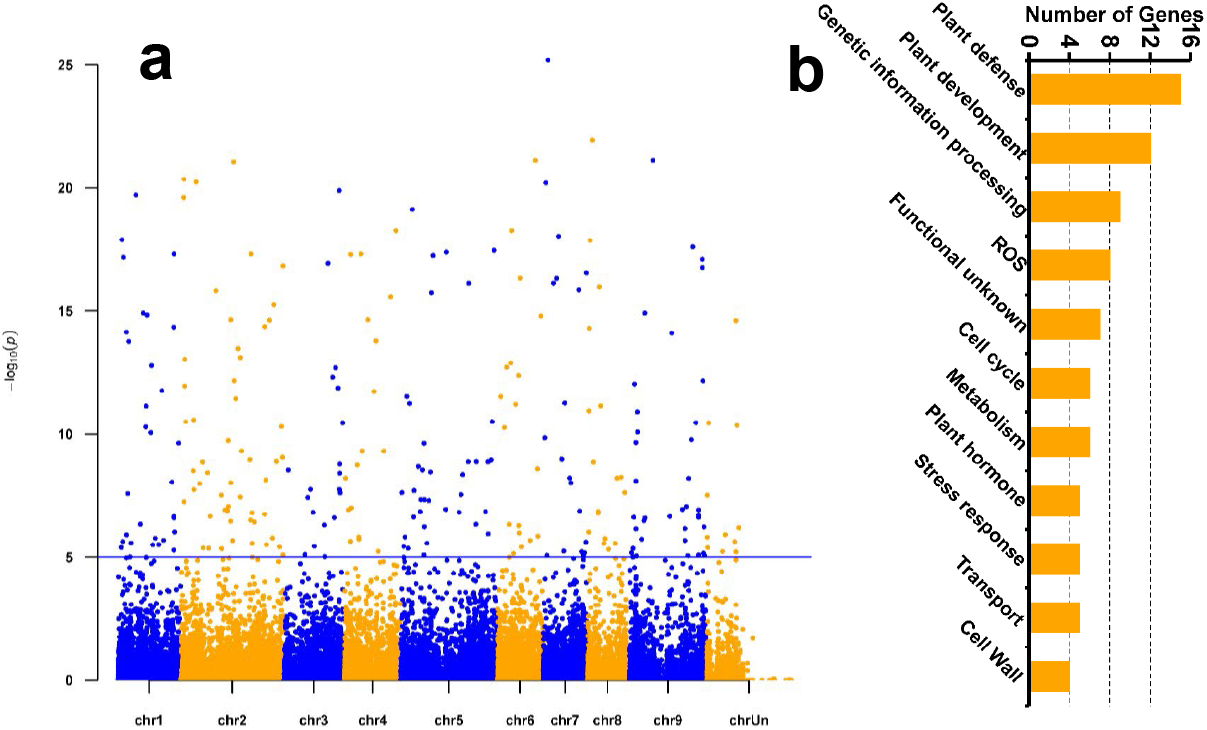
The genomic variations showing significant associations with citrus HLB based on GWAS. Manhattan plots depicting HLB (a) showing significant associations with citrus genomic variations. Points over the blue line represent SNPs showing significant associations with HLB (corrected P-value < 1e-5). b. The functional pathway distribution of genes that contain the HLB associated SNPs.

**Figure 5.**
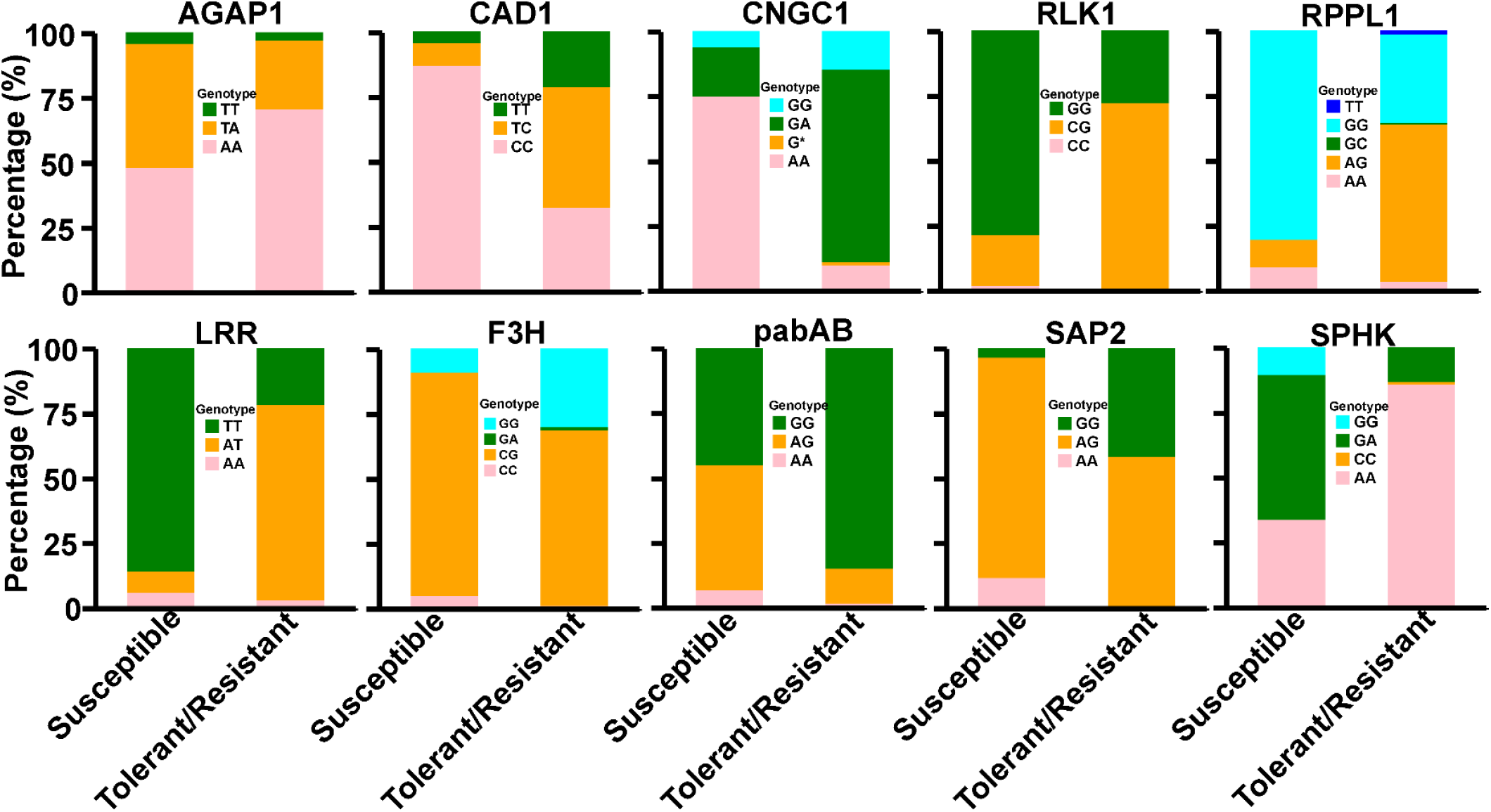
The type of citrus HLB disease associated SNPs among different citrus varieties. These SNPs were from genes involved in plant defense (AGAP1, LLR4, MIK2-LIKE, LRR2, RPPL1 and SUT1), and stress response (SEC11 and SPHK). AGAP1, ACYLATED GALACTOLIPID- ASSOCIATED PHOSPHOLIPASE 1; LLR4, MALECTIN-LIKE DOMAIN (MLD)- AND LEUCINE-RICH REPEAT (LRR)-CONTAINING PROTEIN 4; MIK2-LIKE, MDIS1-INTERACTING RECEPTOR LIKE KINASE2 like; LRR2, Leucine-rich repeat protein 2; RPPL1, disease resistance RPP13-like protein 1; SUT1, SUPPRESSORS OF TOPP4-1; SEC11, signal peptidase I; SPHK, sphingosine kinase.

### Allele-specific expression (ASE)

Next, we used the phased diploid genome assembly of *C. sinensis* to investigate whether ASE contributes to the differences in HLB response between *C. sinensis* ‘Valencia’, a HLB susceptible cultivar, and Sugar Belle mandarin LB9-9, a HLB tolerant cultivar. Both cultivars mainly contain genes originated from either mandarin or pummelo. Mandarin LB9-9 is tolerant to HLB, whereas Valencia sweet orange is susceptible to HLB (Deng *et al*., 2019). We conducted ASE analyses for two set of RNA-seq data of *C. sinensis* ‘Valencia’ and Sugar Belle mandarin LB9-9 including both HLB symptomatic and asymptomatic samples (Ribeiro et al. 2022). A total of 1623 and 1668 genes with ASE were identified for Valencia and LB9-9, respectively (Dataset 3). For Valencia, expression of 892 genes of the mandarin allele was higher than the allele with pummelo ancestry, with 731 displaying the reverse pattern with higher expression of the pummelo allele. For LB9-9, 1012 genes of mandarin origin had higher expression whereas 656 genes of pummelo origin had higher expression (Dataset 3). We further compared the ASE genes and differentially expressed genes (DEGs) between HLB symptomatic vs. asymptomatic Valencia or LB9-9. Intriguingly, 615 DEGs in symptomatic vs asymptomatic Valencia and 484 DEGs of LB9-9 were also ASE genes. Only 177 genes were shared between the DEG and ASE analysis including 166 genes showing similar response to HLB in both Valencia and LB9-9 and only 11 genes showed opposite expression pattern to HLB (Dataset 3). The 11 genes included orange1.1t03406 (RLK), Cs9g12460 (raffinose synthase), orange1.1t03953 (haloacid dehalogenase-like hydrolase domain-containing protein), Cs5g18710 (licodione synthase), Cs5g21200 (indole-3-acetate beta-glucosyltransferase), Cs4g03330 (mandrin 4-coumarate—CoA ligase 1), Cs2g19490 (leucine-rich repeat, cysteine-containing type RLK), Cs2g19440 (Glucose-6-phosphate 1-dehydrogenase), Cs2g04330 (LRR-VIII-2 RLK), Cs2g01740 (peptidase aspartic), and Cs1g12660 (caffeoyl-CoA O-methyltransferase).

## Discussion

We have completed the phased diploid genome assembly of *C. sinensis* ‘Newhall’. There are currently nine assembled genomes for citrus and relatives (Peng *et al*., 2020; Wang *et al*., 2018a; Wang *et al*., 2017; Wu *et al*., 2014; Xu *et al*., 2013; Zhu *et al*., 2019). Even though citrus and its relatives are diploid, none of the nine assembled genomes were chromosome-level phased genomes. A phased genome assembly with allelic information is critical for dissecting the special genetic characteristics, accurately evaluating somatic mutation calling and gene expression, as well as conducting allelic level analysis (Wu et al., 2022). The first chromosome-level phased genome of *C. sinensis* was completed for Valencia sweet orange (Wu *et al*., 2022). ‘Newhall’ navel is also sweet orange but with distinct characteristics from Valencia sweet orange, including the harvest time, navel, seeds, and juice contents. Completion of the chromosome-level phased genome for both Newhall and Valencia will provide useful resource to investigate the underlying genetic determinants for such traits. In addition, Newhall chromosome-level phased genome was successfully used in this study to identify multiple candidate genes contributing to the difference in HLB tolerance between mandarin LB9-9 and Valencia.

We have sequenced 91 citrus accessions in this study. In total, 447 citrus accessions and relatives have been sequenced so far. Importantly, the power of GWAS is boosted when sample size is large (Korte and Farlow, 2013). Imai and colleagues have demonstrated GWAS is suitable for citrus analysis based on analysis of 110 citrus accessions composed of landraces, modern cultivars, and inhouse breeding lines (Imai et al., 2018). The large amount of whole genome sequencing data of citrus and relatives enabled GWAS analysis of putative genetic determinants of HLB resistance/tolerance and susceptibility. Mattia and colleagues conducted GWAS analysis of genes responsible for flavonoid biosynthesis of diverse mandarin accessions (Mattia et al., 2022). Another study by Minamikawa *et al*. used GWAS analysis to identify putative genetic traits controlling fruit quality (Minamikawa et al., 2017). In both studies, SNP arrays were used. With the advances of high-throughput sequencing, whole genome sequencing has become a viable genotyping technology for use in GWAS analyses, offering the potential to analyze a broader range of genome-wide variations (McMahon *et al*., 2021). Whole genome sequence provides a significant advantage over array-based methods, with the potential to detect and genotype all variants present in a sample, not only those present on an array or imputation reference panel (McMahon *et al*., 2021). However, few GWAS analyses have been performed for citrus using whole genome sequencing data to date, probably owing to the challenges in handling a large amount of genomic data. This GWAS analysis using whole genome sequencing data in this study has advanced our understanding of HLB resistance, tolerance or susceptibility in citrus accessions.

Citrus genomic resources generated in the past and in this study enabled identification of candidate genes underlying the HLB resistance, tolerance, or susceptibility via Pan genome analysis of genes presence and absence in 10 citrus accessions, indel analyses of 26 citrus accessions, GWAS analysis of 447 citrus accessions, and ASE analysis of Valencia and mandarin LB9-9. Among the identified genes, NBS-LRR genes are the most abundant. This probably evolves from that CLas resides inside the phloem sieve elements (**Bové**, 2006) and NBS-LRRs are involved in direct recognition of CLas proteins either on the surface or released (Clark et al., 2018; Pang et al., 2020; Prasad et al., 2016). More unique NLRs genes were found in HLB-susceptible citrus accessions than in HLB-tolerant accessions, suggesting CLas might trigger more severe immune responses in HLB susceptible citrus genotypes. More indel mutations were identified in NLR genes in the HLB resistant group than in the susceptible group, which suggest that such mutations might lead to reduced immune responses to CLas in the HLB resistant group. In addition to NBS-LRR, genes encoding CDPK, PRRs, and RLCKs were also identified. CDPKs play critical roles in plant immunity, including regulation of oxidative burst, gene expression, and hormone signal transduction (Liese and Romeis, 2013). PRRs including receptor-like kinases (RLKs) and receptor-like proteins (RLPs) usually localize on the membrane to detect MAMPs in the apoplast. It is possible that some PAMPs of CLas, such as LPS and flagella (Andrade et al., 2020; Zou et al., 2012), are released into the apoplast to be sensed by citrus cells. RLCKs including BOTRYTIS-INDUCED KINASE 1 (BIK1) and related PBS1-like kinases act synergistically with multiple PRRs to allow subsequent phosphorylation of the transmembrane NADPH-oxidase RESPIRATORY BURST OXIDASE D (RBOHD), the main producer of ROS during pathogen infection (Couto et al., 2016; Kadota et al., 2014; Liang et al., 2016). Consistent with our observation, it was reported that CLas causes less expression changes in immunity genes for the HLB-tolerant US-897 than the HLB-susceptible ‘Cleopatra’ mandarin (Albrecht and Bowman, 2012), which might result from the differences in the NLR, RLK, RLCK, and CDPK genes. Intriguingly, 74 antioxidant biosynthesis or antioxidant enzyme genes were absent in susceptible citrus accessions, but were present in one or more tolerant citrus accessions. This is consistent with that higher antioxidant levels and antioxidant enzyme activities were reported to account for the higher tolerance of Persian triploid lime than Mexican lime (Sivager *et al*., 2021). Since the presence/absence pattern of antioxidant biosynthesis or antioxidant enzyme genes was not universal, they are likely responsible for some citrus genotypes, but not all genotypes. For example, HLB-tolerant Sugar Belle mandarin LB9-9 has similar antioxidant enzyme activities as HLB-susceptible Valencia sweet orange, but has higher phloem regeneration capacity than the later. Antimicrobial peptides were also reported to be responsible for HLB resistance in Australian Finger lime (Huang *et al*., 2021).

Overall, this study has provided a phased chromosome-level genome assembly for *C. sinensis* ‘Newhall’, and sequenced 91 new citrus accession, which were used for identification of putative genes responsible for HLB resistance, tolerance, or susceptibility. Such genes should be further verified using other approaches such as mutagenesis or gene silencing. Identification of genetic determinants responsible for HLB resistance, tolerance, or susceptibility in citrus accessions provide useful targets for developing of HLB-resistant/tolerant citrus cultivars using the CRISPR genome editing tool (Huang et al., 2022; Jia et al., 2022).

## Methods

### Plant materials

The Newhall navel sweet orange (*C. sinensis* Osbeck cv. Newhall) was sequenced in this study. The plant materials of Newhall navel sweet orange were obtained for genomic DNA and RNA extractions from the National Navel Orange Engineering Research Center, Ganzhou, Jiangxi Province, China. Leaf and root samples of citrus germplasms were collected from the Lower Variety Grove, California Citrus State Historic Park, Riverside, California for GWAS study.

### DNA and RNA extraction

Genomic DNA for Illumina sequencing was extracted using the phenol-chloroform method (Rana et al., 2019). Genomic DNA was isolated using Nanobind Plant Nuclei Big DNA Kit (Circulomics Inc., Baltimore, MD, USA) following the manufacturer’s instructions for PacBio and Hi-C sequencing. For full length transcript sequencing (Iso-Seq), the RNA was extracted using an RNAprep Plant Kit (Qiagen, Valencia, CA, USA). Genomic DNA for BGI sequencing was extracted using MoBio Powersoil DNA extraction kit (MoBio Laboratories Inc. Carlsbad, CA, USA) following the manufacturer’s instructions. The quality of genomic DNA and RNA was evaluated using agarose gel electrophoresis and Qubit 3.0 Fluorometer (Life Technologies, USA).

### Library construction and sequencing

Following the manufacturer’s protocol of short read DNA sequencing from Illumina (Kozarewa et al., 2009), the library was prepared. After quality control, quantification, and normalization of the DNA libraries, 150-bp paired-end reads were generated using the Illumina NovaSeq 6000 platform according to the manufacturer’s instructions.

PacBio HiFi SMRTbell Library with 15 kb DNA fragment was constructed following the manufacturer’s protocol. The constructed library was then sequenced by Pacbio Sequel II platform according to the manufacturer’s instructions.

Hi-C libraries were prepared by a standard procedure (Belton et al., 2012). Formaldehyde solution was used to fix plant cells. Then, 2.5 M glycine was added to quench the cross-linking reaction. The crosslink DNA was treated with restriction enzymes (e.g., *Hin*dIII), creating a gap on both sides of the crosslinking point. The exposed DNA ends were repaired and covalently linked with biotin-14-dCTP (Invitrogen Life Technologies, Carlsbad, CA), and then connected by T4 DNA ligase (Invitrogen Life Technologies, Carlsbad, CA) to form a closed random circular DNA structure. Proteinase K (Invitrogen Life Technologies, Carlsbad, CA) was used to digest proteins at the junction point to disconnect the crosslink of the protein and DNA. Genomic DNA was extracted and later fragmented into 350 bp. Capturing and enriching biotin-labeled fragments was conducted via the affinity of streptomycin to biotin, which was used to construct Illumina library. Hi-C sequencing libraries were amplified by PCR and sequenced on Illumina HiSeq-2500 platform (PE 125 bp). The 90 citrus genomes for GWAS study were sequenced using BGISEQ500 platform. Shotgun genomic library preparation and sequencing were performed per the manufacturer’s protocol at BGI-Shenzhen, China. Briefly, 500 ng of input DNA was used for library generation and fragmented ultrasonically to yield 400 to 600 bp of fragments. DNA fragments were then end-repaired and A-tailed, and adaptors with specific barcodes were added. PCR amplification of DNA fragments was carried out to generate a single-strand circular DNA library. The DNA libraries were sequenced by BGISEQ500 using a paired-end 100-bp sequencing strategy. On average, more than 41.9 Gb of raw data were generated for each genomic sample for GWAS analysis.

### De novo genome assembly for Newhall navel sweet orange

Hifiasm v 0.15.4 (Cheng et al., 2021) with default parameters was used to assemble HiFi reads into scaffolds, which were corrected using short Illumina DNA reads. Based on Hi-C sequencing data, the assembled scaffold sequences were mounted to the near-chromosome level using ALLHIC v 0.9.8 (Zhang et al., 2019). According to chromosome interaction intensity using juicebox v1.11.08 (Durand et al., 2016), the near-chromosome level genome was manually corrected into a chromosome-level genome.

### Genome quality assessment

To assess the assembly quality of the assembled genomes, the completeness of the assembly was evaluated using Benchmarking Universal Single-Copy Orthologs (BUSCO) v10 (Manni et al., 2021), and the Conserved Core Eukaryotic Gene Mapping Approach (CEGMA) (Parra et al., 2007). The coverage of assembled genomes was also calculated by mapping Illumina short reads to the assembly using Burrows-Wheeler Aligner (BWA) (Li and Durbin, 2010).

### Repeat element identification

Repeat elements of whole genome were identified using both homology alignment and de novo prediction. Tandem repeat was extracted using TRF (Benson, 1999) by ab initio prediction. The de novo repetitive element library was constructed using LTR_FINDER (Xu and Wang, 2007), RepeatScout (Lian et al., 2016), and RepeatModeler (Flynn et al., 2020). Homolog repeats were predicted based on Repbase database (Jurka et al., 2005) employing RepeatMasker software (Tempel, 2012) and its in-house scripts RepeatProteinMask with default parameters. Using uclust program, a non-redundant transposable element (TE) library (a combination of homolog repeats and de novo TE) was generated, which was applied to mask the genome using RepeatMasker software.

### Gene structure annotation

Homology-based prediction, ab initio prediction, and RNA-Seq assisted prediction were employed to perform the gene model prediction. Genomic sequences were aligned to homologous proteins using tblastn v2.2.26 (Camacho et al., 2009) with a threshold of E-value ⩽ 1e−5. Based on the matched proteins from reference genomes, GeneWise (v2.4.1) (Birney et al., 2004) software was used to predict gene structure. Augustus v3.2.3 (Nachtweide and Stanke, 2019), Geneid v1.4 (Alioto et al., 2018), Genescan v1.0 (Burge and Karlin, 1997), GlimmerHMM v3.04 (Majoros et al., 2004), and SNAP_2013-11-29 were used for the automated de novo gene prediction. The transcriptome assembly was performed using Trinity v2. 1. 1 (Grabherr et al., 2011; Haas et al., 2013) for the genome annotation. To identify exon region and splice positions, the RNA-Seq reads from leaf, root, and fruit tissues were aligned to genome using Hisat v2.0.4 (Kim et al., 2019) with default parameters. The alignment results were then used as input for StringTie v1.3.3 (Pertea et al., 2015) with default parameters for genome-based transcript assembly. The non-redundant reference gene set was generated by merging genes predicted from three methods using EvidenceModeler (EVM) v1. 1. 1 (Haas et al., 2008).

### Gene functional annotation

Gene functions were assigned according to the best match by aligning the protein sequences to the Swiss-Prot (Boutet et al., 2007) using Blastp (Camacho *et al*., 2009) with a threshold of E-value ⩽ 1e−5 (Gertz et al., 2006). The motifs and domains were annotated using InterProScan70 v5.31 (Quevillon et al., 2005) by searching publicly available databases, including ProDom (Servant et al., 2002), PRINTS (Attwood et al., 1994), Pfam (Mistry et al., 2021), SMRT (Lou et al., 2020), PANTHER (Mi et al., 2013; Mi et al., 2019) and PROSITE (Hulo et al., 2006). The Gene Ontology (GO) (Consortium, 2015) IDs for each gene were assigned according to the corresponding InterPro (Hunter et al., 2009) entry. We predicted the proteins function by transferring annotations from the closest BLAST hit with a threshold of E-value <10^−5^ from the Swiss-Prot (Boutet *et al*., 2007) and the NR databases (Pruitt et al., 2007) using DIAMOND v0.8.22 (Buchfink et al., 2015). We also mapped gene set to the KEGG (Kanehisa et al., 2017) and eggNOG databases to generate more comprehensive gene functional information. To infer the evolutionary trajectories of genes in Newhall navel orange, we calculated the number of substitutions per synonymous site (Ks) and the number of substitutions per nonsynonymous site (Ka) for each homologous gene pairs. A Ka/Ks ratio more than 1, less than 1, and equal to 1 indicates positive selection, purifying selection and neutral evolution, respectively (Gao et al., 2017; Lynch and Conery, 2000).

### Non-coding RNA annotation

tRNAs were predicted using the program tRNAscan-SE (Lowe and Chan, 2016). rRNA sequences were predicted using Blast (Camacho *et al*., 2009) with relative species’ rRNA sequences. Other ncRNAs, including miRNAs and snRNAs, were identified by searching the Rfam database (Kalvari et al., 2021) with default parameters.

### Identification of genomic variations

We used For SNPs and InDels identification, short sequencing reads were aligned to the high-quality sweet orange genome as the reference genome (Wang et al., 2021) using BWA software. Duplicated mapping reads and unmapped reads were removed with the SAMtools (Li et al., 2009). All genotype information for the polymorphic sites was generated using the GATK population method (McKenna et al., 2010). The generated SNPs and InDels were filtered by quality and sequence depth.

### Pan genome construction

To construct the citrus pan genome, we generated the gene families from 10 citrus and relative genomes (Table S1). For each genome, a gene containing CDS with 100% similarity to other genes was removed by using the cd-hit-est of CD-HIT v4.8 toolkit (Li and Godzik, 2006). Protein sequences of the remaining genes were subjected to homologous searching by BLASTp (Camacho *et al*., 2009) with default parameters. OrthoFinder v2.2.7 (Emms and Kelly, 2019) was used to deal with the BLAST result with default parameters to make gene family clustering. The gene families shared among all accessions were defined as core gene families. The gene families that were missed in one or two accessions were defined as softcore gene families. The gene families that were missed in more than two accessions were defined as dispensable gene families, and gene families that only existed in one accession were defined as private gene families. We used eggNOG database to generate gene functional information for the citrus pan genome.

### GWAS

To generate comprehensive SNP genotyping data for GWAS analysis, we sequenced 91 new citrus genomes and collected 356 representative genomic data of citrus and its relatives from public database (Table S20, Table S21). Raw reads for each genomic sample were filtered by SOAPnuke (v1.5.3) with the parameters set as “filterMeta -Q 2 -S -L 15 -N 3 –P 0.5 -q 20 -l 60 -R 0.5 -5 0” (Chen et al., 2018). The trimmed reads were mapped to the sweet orange genomes (Wang *et al*., 2021; Xu *et al*., 2013), *C. clementina* (Wu *et al*., 2014), pummelo, Citron, kumquat, Atlantia, Papeda (Wang *et al*., 2017), *C. mangshanensis*(Wang *et al*., 2018a), and Swingle citrumelo (Zhang et al., 2016) genomes using Bowtie2 software (Langmead and Salzberg, 2012) to identify the citrus reads. The high quality paired-end short citrus genomic reads were mapped to pummelo (*C. maxima*) (Wang *et al*., 2017) reference genome, which is of the highest-quality among sequenced genomes, using SOAP. Duplicated mapping reads were removed with the SAMtools package (Li *et al*., 2009). All genotype information for the polymorphic sites was generated using the GATK population method (McKenna *et al*., 2010). The generated SNPs were filtered by quality and sequence depth, and SNPs with minor allele frequency (MAF) more than 0.01 and individual level missingness less than 10% were retained for GWAS analysis.

To perform the association between HLB and citrus genomic variations, we collected the HLB resistant or tolerant information for each citrus genotypes from previous studies (Table S20). The population structure of citrus was estimated using PCA method (Minamikawa *et al*., 2017). GWAS for HLB was conducted using the univariate linear mixed model in the GEMMA package, which accounts for population stratification using the first two principal components (PCs) of population structure and kinship matrices (Zhou and Stephens, 2014). P-values for multiple testing were corrected using the FDR method (Storey, 2003). All items with corrected P-values below 1e-5 were considered significant. Significantly associated SNPs were mapped to the citrus reference genome to acquire gene annotations. The genes containing significantly associated SNPs were functionally annotated using the KEGG and agriGO 2.0 databases (Tian et al., 2017).

### Allelic Specific Expression of transcriptomes of Valencia sweet orange and Mandarin LB9-9 in response to CLas infection

We generated the allelic specific expression of genes (ASE) of Valencia sweet orange and Mandarin LB9-9 based on the method described previously (Sun *et al*., 2020). Cleaned RNA-seq reads were first mapped against both alleles of genome of Newhall navel sweet orange using HISAT2 (Kim *et al*., 2019) and SAMtools (Li *et al*., 2009) to select good-quality alignments. Reads uniquely aligned to the same chromosome in both alleles were preserved and assigned to the allele if the alignment had a higher score and fewer mismatches than to the other allele. Reads that failed to be assigned, which were mainly from homozygous genomic region or regions that were unassembled or haplotype unresolved, were not considered in downstream analyses. The allele-specific reads were mapped to the Newhall navel sweet orange consensus genome using HISAT2 and SAMtools. Raw counts for each genes were quantified using HTSeq-count (Anders et al., 2015). Genes with total counts >10 in all biological replicates were analyzed for ASE using DESeq2 (Love et al., 2014). A gene was considered to have ASE if the expression difference of the two alleles was significantly greater than twofold (adjusted P < 0.05).

## Figure legends

Dataset 1. Immunity gene presence and absence analysis in 10 HLB-susceptible and -tolerant citrus accessions

Dataset 2. The information of SNP that significantly associated with citrus HLB disease

Dataset 3. Allelic specific expression analysis of Valencia sweet orange and Mandarin LB9-9 in response to CLas infection

