## Supplementary Information for "Citrus genomic resources unravel putative genetic determinants of Huanglongbing, a pathogen-triggered immune disease"

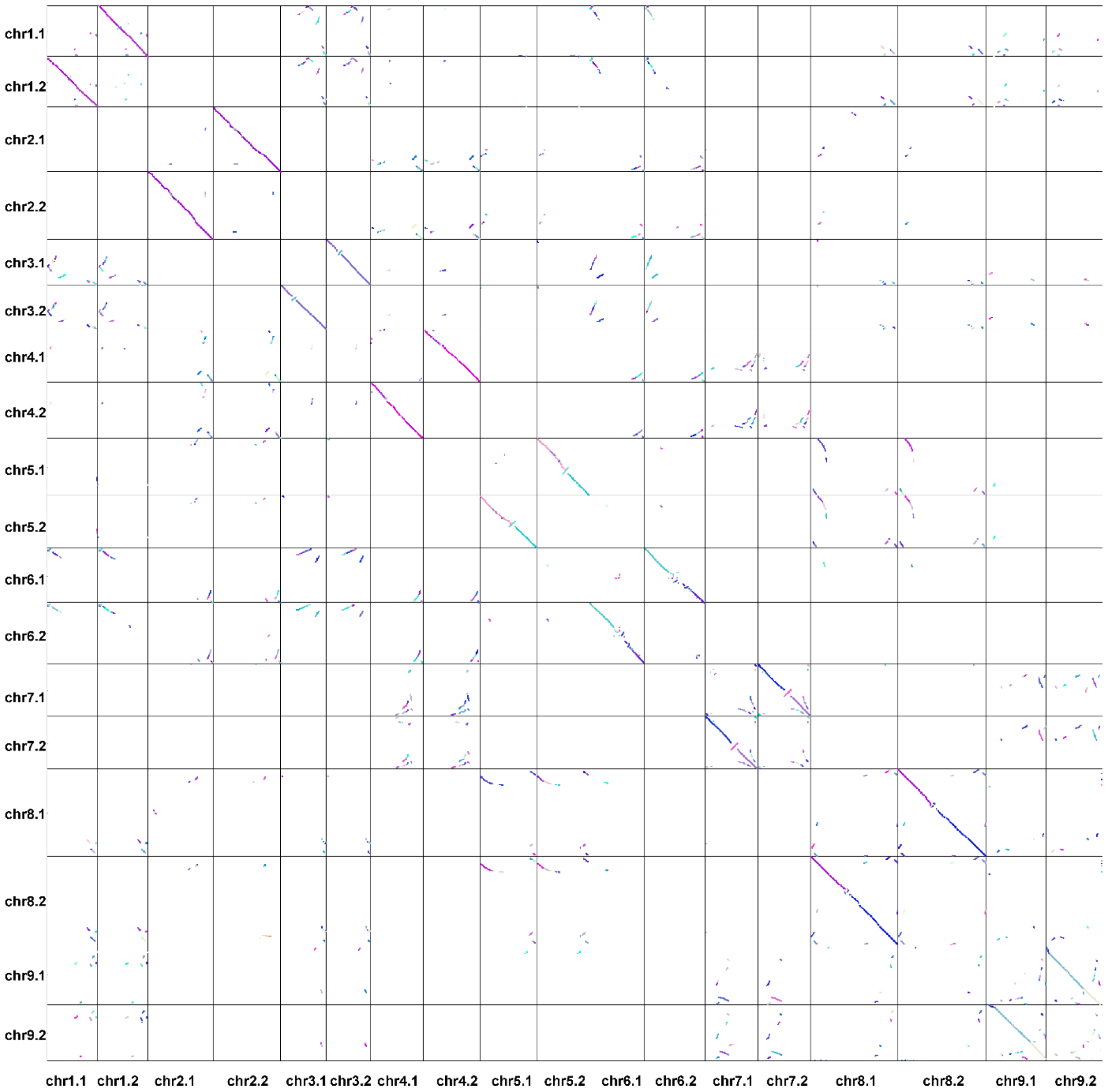


**Figure S1.** Synteny between homologous chromosomes of diploid Newhall navel orange genome.

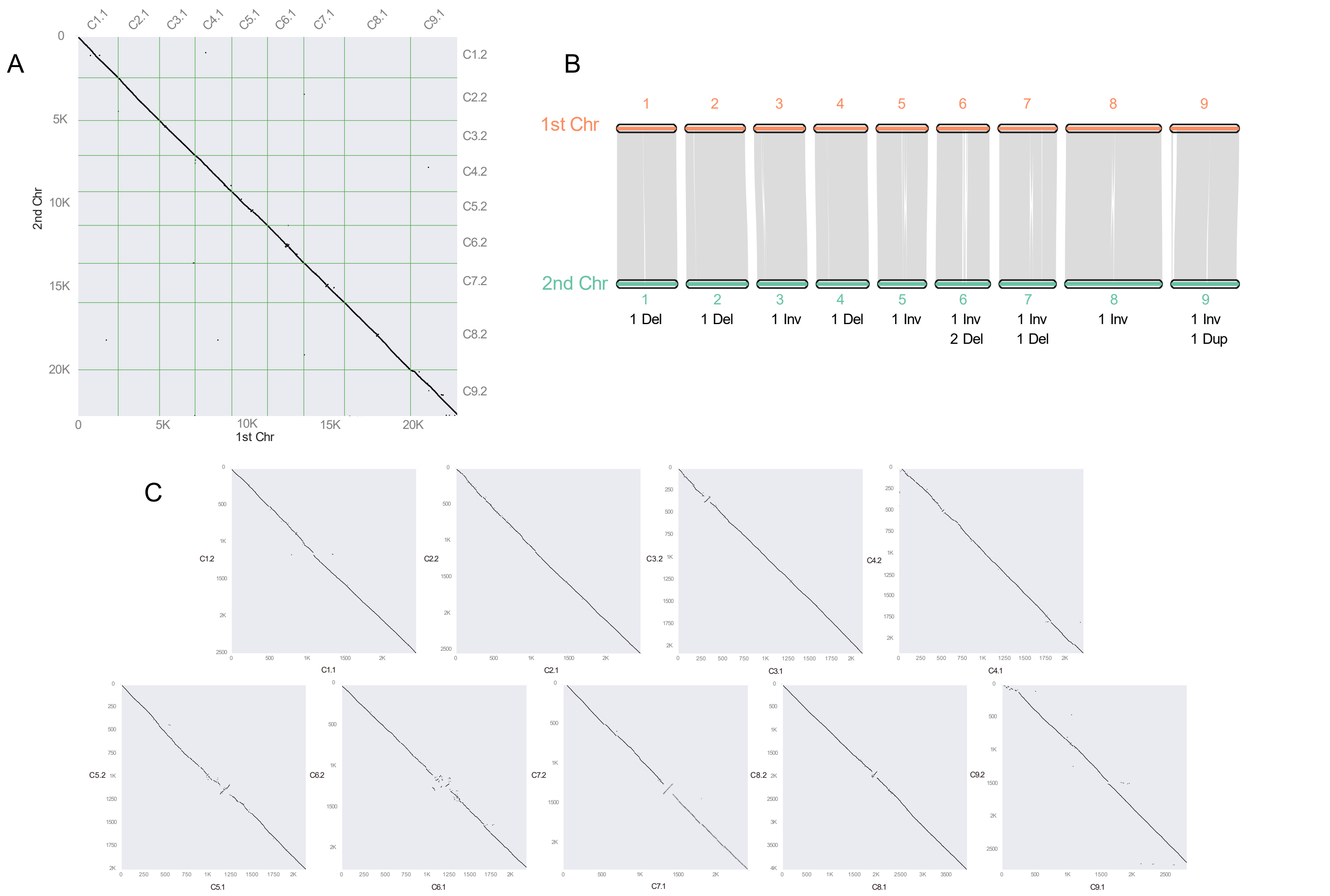


**Figure S2.** Synteny block of chromosomes of Newhall navel orange genome. A. Synteny between homologous chromosomes. B. Structural variations between homologous chromosomes (Del: deletions/insertions, Inv: inversions, Dup: duplication). C. Synteny between homologous sequences for each paired chromosome.

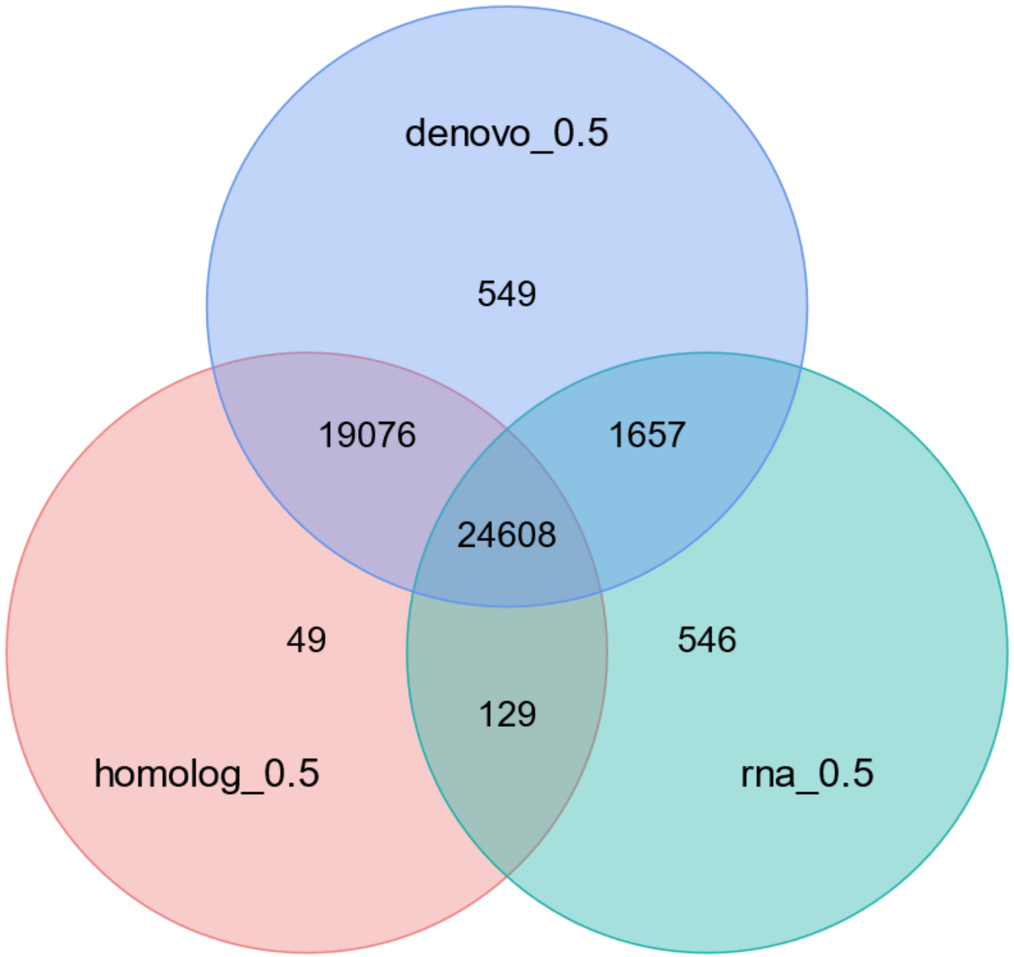


**Figure S3**. Gene predication in Newhall navel orange genome. De novo: the genes supported by the De novo prediction in EVM. Homolog: homology-based prediction. RNA: the genes supported by the RNA-seq in prediction in EVM. Each evidence support was based on gene overlap greater than 50%. Numbers indicate the number of genes.

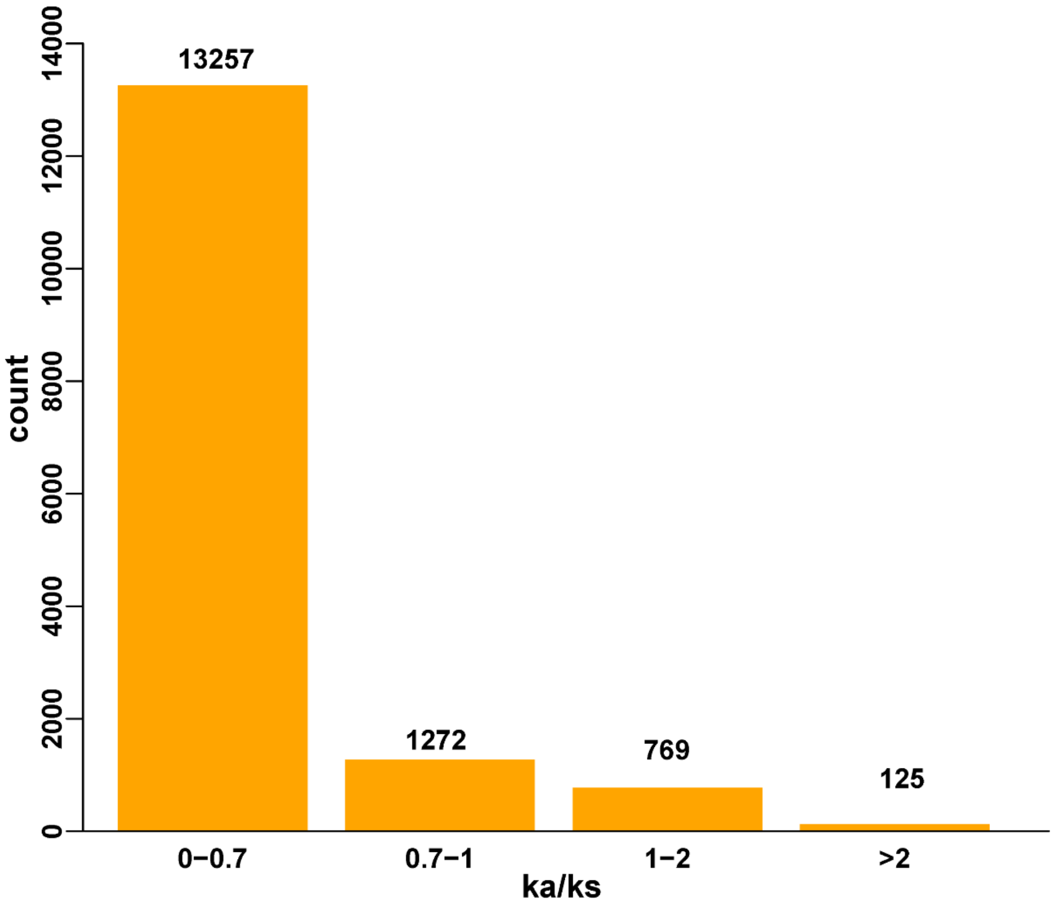

**Figure S4.** Histogram of the Ka/Ks of homologous genes in Newhall navel orange genome. The horizontal axis indicates the range of ka/ks values, and the vertical axis indicates the gene count. Ka/Ks: The ratio of the number of nonsynonymous substitutions per nonsynonymous site (Ka) to the number of synonymous substitutions per synonymous site (Ks).

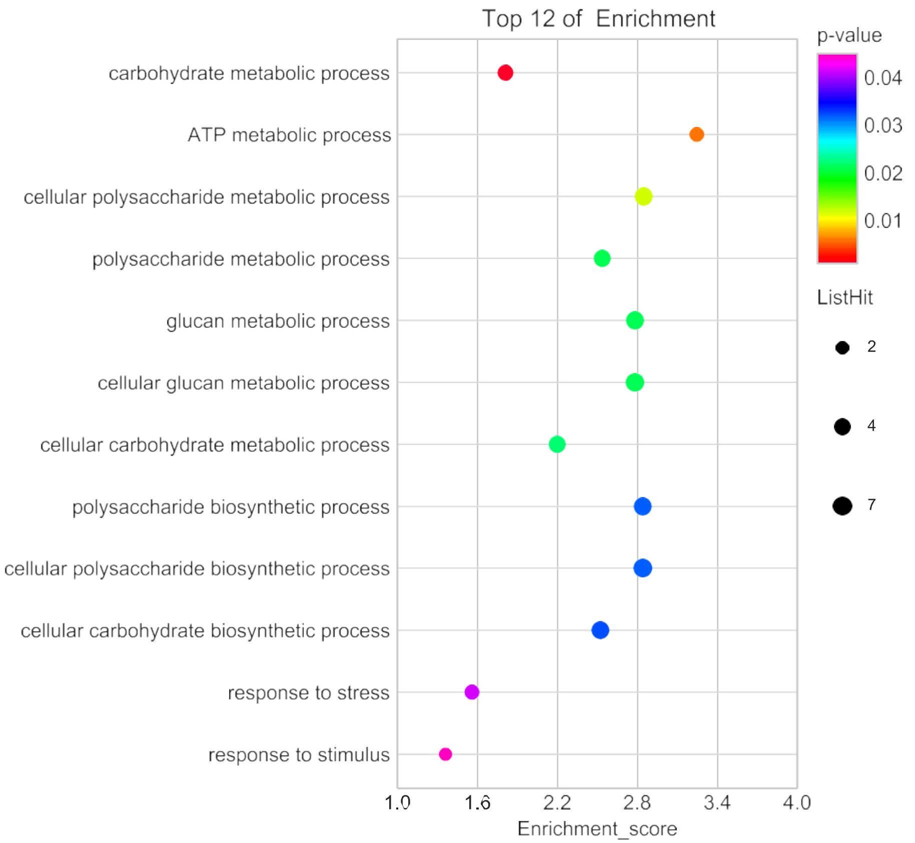


**Figure S5.** The GO annotation of genes underwent positive selection.

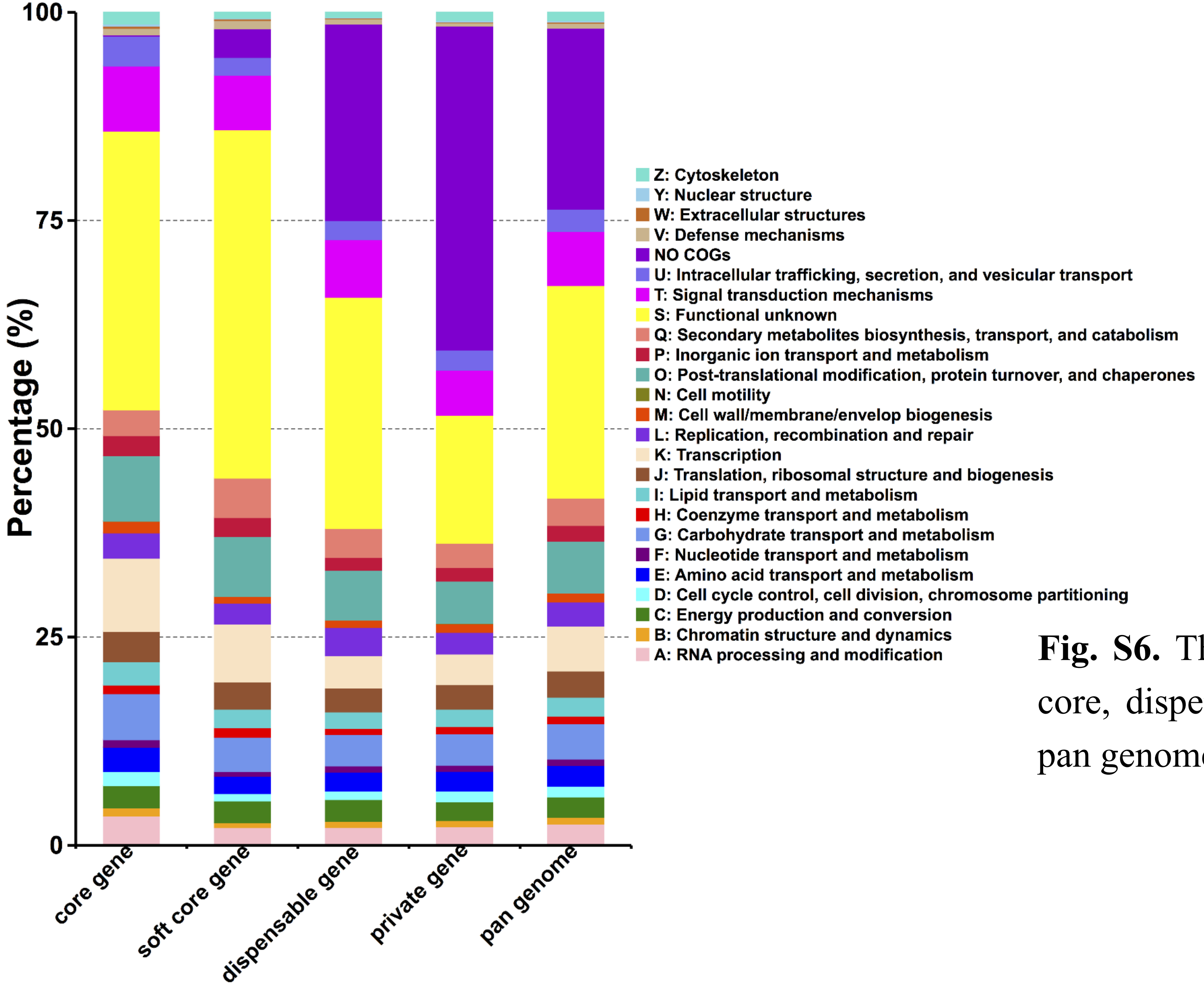


**Fig. S6.** The COG distribution for core, soft core, dispensable and private genes in citrus pan genome.

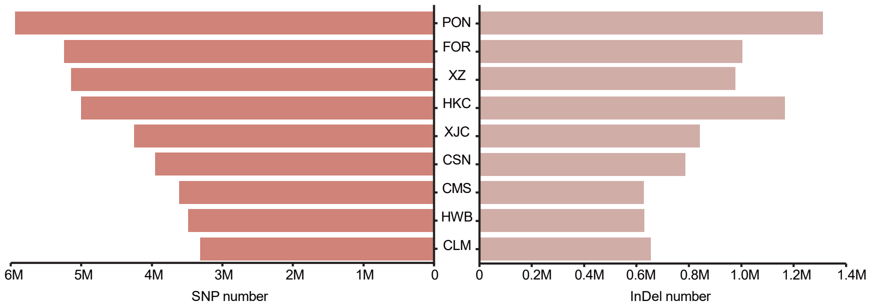


**Figure S7.** The genomic variations of 9 accessions of citrus and relatives. The left side is the number of SNP (single-nucleotide polymorphism) for each genome. The right side is the number of InDels for each genome. (HKC: *Atlantia buxifolia*, CSV: ‘Valencia’ sweet orange, FOR: *Fortunella hindsii*, CSN: ‘Newhall’ sweet orange, XZ: *Citrus medica*, XJC: *Citrus ichangensis*, HWB: pummelo, CMS: *Citrus reticulata* 'Mangshan', PON: *Poncirus trifoliata*, and CLM: Clementine mandarin).

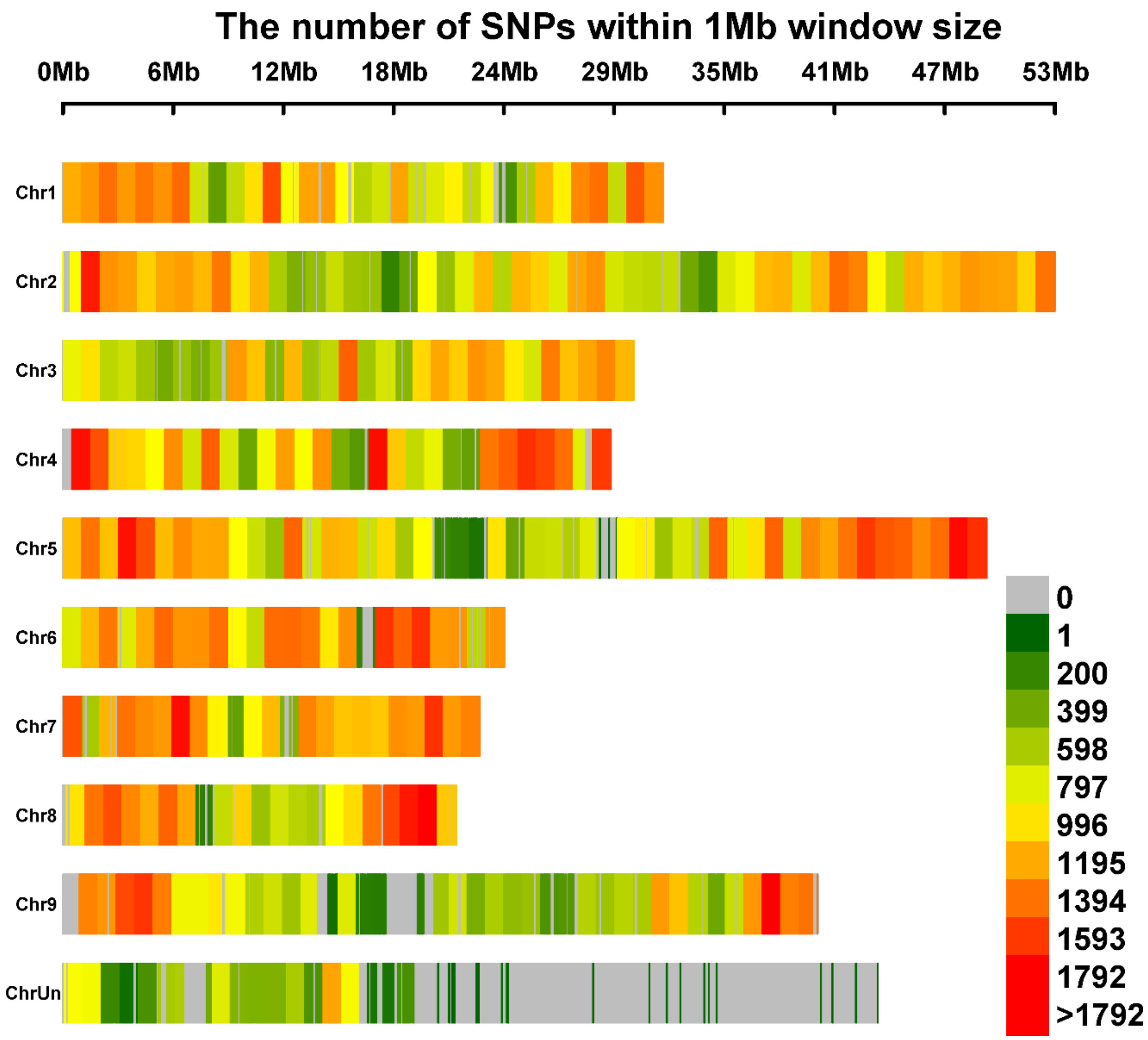


**Figure S8.** The SNPs of 447 accessions of citrus and relatives that are associated with HLB by GWAS analysis.

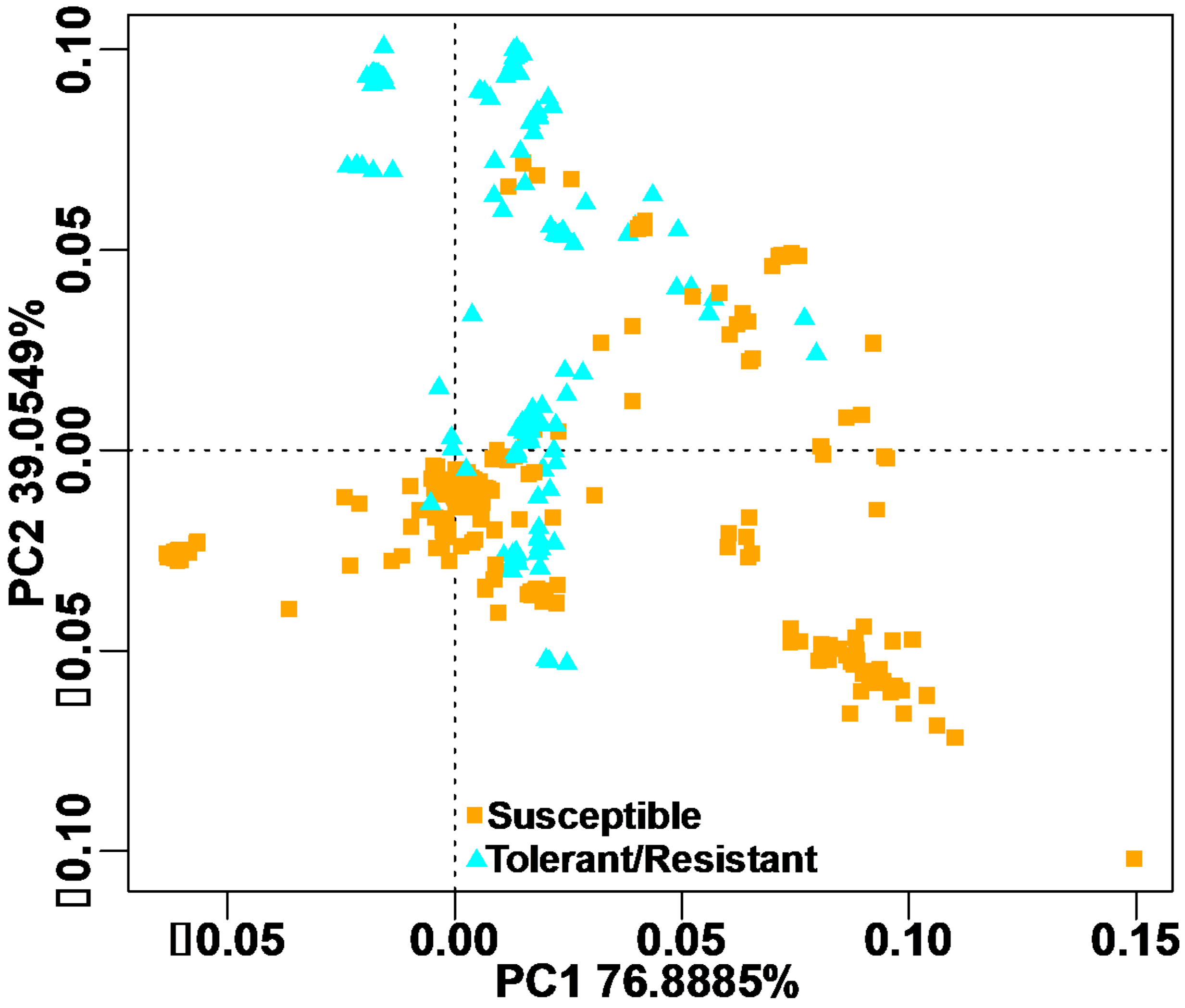


**Figure S9.** The population structure of 447 accessions of citrus using in GWAS analysis.

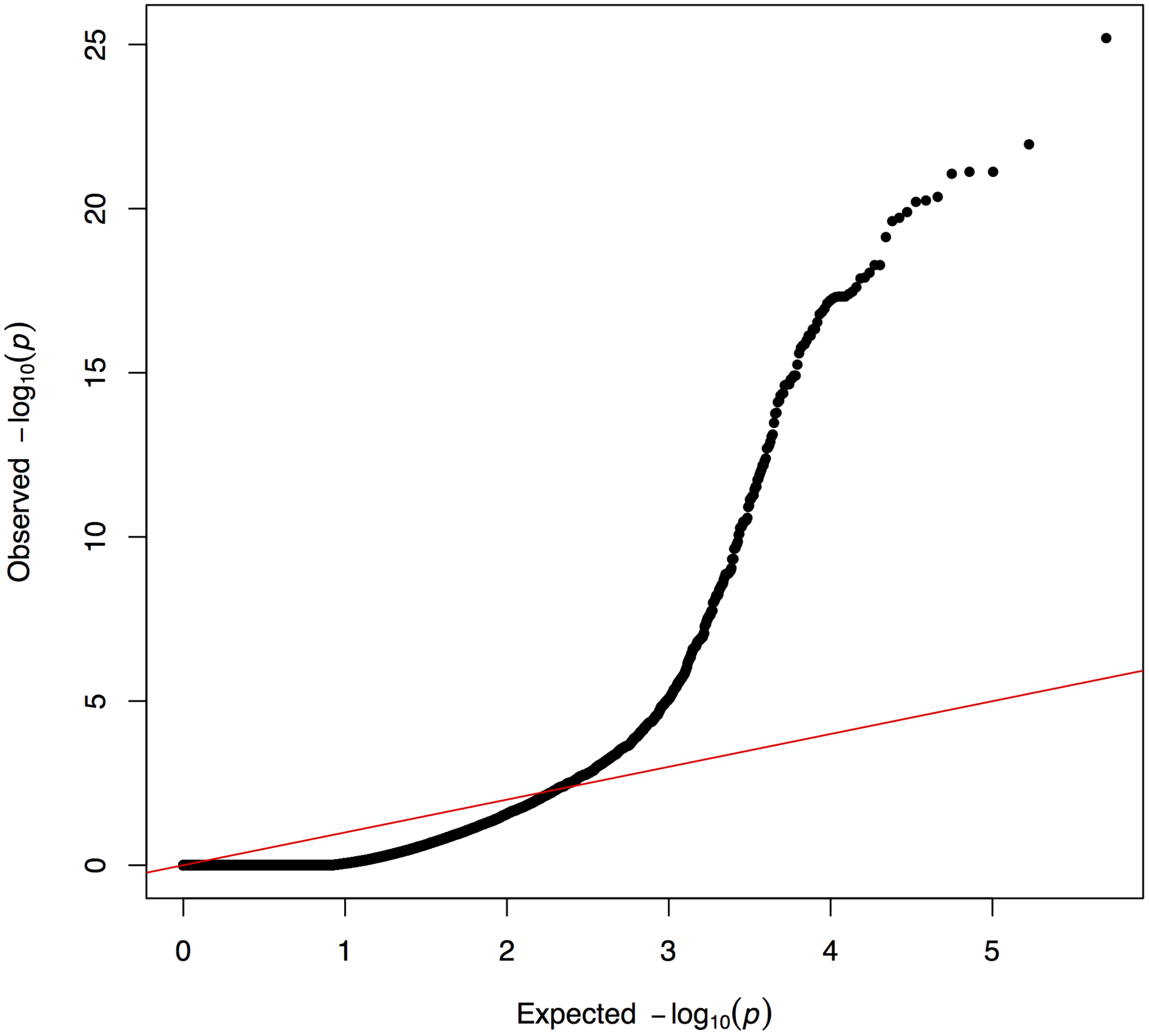


**Figure S10.** The Quantile-Quantile Plot of GWAS analysis for 447 accessions of citrus and relatives that are associated with HLB.

**Table S1. Sequencing data summary of Newhall navel orange**

| Sequencing technology | Insert size | Total data (G) | Read length (bp) | Sequence coverage (X) |
|---|---|---|---|---|
| Illumina reads | 350 bp | 54 | 150 | 77.14 |
| PacBio reads | -- | 137 | -- | 195.63 |
| PacBio HIFI | 15 k | 35 | 14,554 | 50 |
| Hi-C | -- | 45 | -- | 64.29 |
| Iso-Seq of leaf | -- | 14 | -- | -- |
| Iso-Seq of root | -- | 21 | -- | -- |

**Table S2. Genome assembly summary of Newhall navel orange**

| Item | Value |
|---|---|
| Estimate of genome size (Mb) | 700 |
| Chromosome number (2n) | 18 |
| Total size of genome sequence (Mb) | 685 |
| Number of contigs (>100bp) | 1,624 |
| N50 length of contigs (Mb) | 12.5 |
| Number of scaffolds | 1,013 |
| N50 length of scaffolds (Mb) | 32.86 |
| GC content | 36.83 |
| Number of gene models | 46,616 |
| Mean transcript length (bp) | 3025 |
| Mean CDS length (bp) | 1175 |
| Mean number of exon per gene | 5 |
| Mean exon length (bp) | 231 |
| Mean intron length (bp) | 454 |
| Number of predicted miRNA gene | 931 |
| Total length of TE (bp) / % in genome | 363,999,943/53.12 |

**Table S3. The ATGC bases distribution of Newhall navel orange genome**

| Base | Number (bp) | Percentage of genome (%) |
|---|---|---|
| G | 126,026,314 | 18.39 |
| T | 216,564,487 | 31.6 |
| C | 126,381,518 | 18.44 |
| A | 216,240,679 | 31.55 |
| N | 61,100 | 0.009 |
| Total | 685,274,098 | -- |
| GC | 252,407,832 | 36.83 |

**Table S4. The length and cluster number for each chromosome in Newhall navel orange genome**

| Chromosome ID | Cluster Number | Sequence Length (bp) |
|---|---|---|
| Chr1.1 | 3 | 29,757,323 |
| Chr1.2 | 29 | 29,428,234 |
| Chr2.1 | 14 | 37,800,815 |
| Chr2.2 | 19 | 39,622,446 |

| | | |
|---|---|---|
| Chr3.1 | 13 | 26,553,055 |
| Chr3.2 | 4 | 25,721,200 |
| Chr4.1 | 50 | 31,191,295 |
| Chr4.2 | 36 | 33,146,896 |
| Chr5.1 | 31 | 33,215,067 |
| Chr5.2 | 20 | 30,814,586 |
| Chr6.1 | 50 | 32,665,665 |
| Chr6.2 | 87 | 36,026,403 |
| Chr7.1 | 75 | 30,969,761 |
| Chr7.2 | 24 | 30,849,564 |
| Chr8.1 | 13 | 50,940,101 |
| Chr8.2 | 22 | 51,415,404 |
| Chr9.1 | 23 | 35,155,523 |
| Chr9.2 | 116 | 32,860,738 |

**Table S5. The haploid genomic features statistics using K-mer (17-mer) method**

| Item | Value |
|---|---|
| Kmer length | 17 |
| Depth | 108 |
| Number of kmer | 38,417,033,382 |
| Genome size (M) | 355.71 |
| Revised Genomesize (M) | 350.15 |
| Heterozygous rate (%) | 1.8 |
| Repeat rate (%) | 47.23 |

**Table S6. The BUSCO assessment of Newhall navel orange genome**

| Item | BUSCO notation assessment results |
|---|---|
| Complete BUSCOs | 95.10% |
| Complete and single-copy BUSCOs | 85.80% |
| Complete Duplicated BUSCOs | 9.30% |
| Fragmented BUSCOs | 1.50% |
| Missing BUSCOs | 3.40% |
| Total BUSCO groups searched | 1440 |

**Table S7. The CEGMA assessment of Newhall navel orange genome**

| Complete | | Complete + Partial | |
|---|---|---|---|
| # Prots | % Completeness | # Prots | % Completeness |
| 226 | 91.13 | 230 | 97.74 |

**Table S8. The sequencing coverage of Newhall navel orange genome**

| Item | | Value |
|---|---|---|
| Reads | Mapping rate (%) | 94.46 |
| Genome | Average sequencing depth | 122.91 |
| | Coverage (%) | 99.97 |
| | Coverage at least 4X (%) | 99.95 |
| | Coverage at least 10X (%) | 99.91 |
| | Coverage at least 20X (%) | 99.83 |

**Table S9. Sequence similarity of syntenic chromosomes**

| Chromosome pair | Length of matched bases (bp) | Alignment block length (bp) | Sequence similarity (%) |
|---|---|---|---|
| chr1.1/chr1.2 | 3,783,651 | 8080977 | 46.82 |
| chr2.1/chr2.2 | 3,499,236 | 8794148 | 39.79 |
| chr3.1/chr3.2 | 3,069,609 | 6954824 | 44.14 |
| chr4.1/chr4.2 | 3,228,645 | 7407252 | 43.58 |
| chr5.1/chr5.2 | 2,880,441 | 6904942 | 41.72 |
| chr6.1/chr6.2 | 2,844,086 | 7601520 | 37.41 |
| chr7.1/chr7.2 | 3,774,356 | 8190481 | 46.08 |
| chr8.1/chr8.2 | 6,021,883 | 14189003 | 42.44 |
| chr9.1/chr9.2 | 3,891,365 | 9164025 | 42.46 |

**Table S10. The statistics of repeat sequences in Newhall navel orange genome**

| Item | Length (bp) | Percentage of genome (%) |
|---|---|---|
| DNA | 11,944,797 | 1.74 |
| LINE | 5,666,545 | 0.83 |
| SINE | 392,380 | 0.06 |
| LTR | 304,403,972 | 44.42 |
| Unknown | 44,112,176 | 6.44 |
| Total | 356,312,523 | 52.00 |

**Table S11. Gene prediction in Newhall navel orange genome**

| Prediction method | | Number of gene | Average transcript length (bp) | Average CDS length (bp) | Average exons per gene | Average exon length (bp) |
|---|---|---|---|---|---|---|
| De novo | Augustus | 46,304 | 2635 | 1169 | 4.9 | 239 |
| | GlimmerHMM | 74,547 | 6521 | 769 | 3.33 | 231 |
| | SNAP | 58,495 | 2662 | 668 | 4.17 | 160 |

| | | | | | | |
|---|---|---|---|---|---|---|
| | Geneid | 58,794 | 4477 | 935 | 4.61 | 203 |
| | Genscan | 9,832 | 13191 | 1402 | 6.85 | 205 |
| Homolog | Csn | 44,682 | 2425 | 1049 | 4.54 | 231 |
| | Cre | 43,770 | 2788 | 1097 | 4.81 | 228 |
| | Cma | 45,768 | 2611 | 1063 | 4.66 | 228 |
| | Csi | 45,017 | 2633 | 1085 | 4.68 | 232 |
| | Cse | 45,144 | 2617 | 1080 | 4.66 | 232 |
| RNA-seq | PASA | 92,740 | 2985 | 1029 | 5.3 | 194 |
| | Transcripts | 38,625 | 4764 | 1875 | 6.34 | 296 |
| EVM | | 52,231 | 2839 | 1102 | 4.76 | 232 |
| Pasa-update | | 51,925 | 2838 | 1116 | 4.79 | 233 |
| Final set | | 46,616 | 3025 | 1175 | 5.08 | 231 |

**Table S12. The statistics of gene functional annotation in Newhall navel orange genome**

| Item | Number | Percentage (%) |
|---|---|---|
| Total | 46616 | -- |
| Swissprot | 35091 | 75.28 |
| Nr | 44599 | 95.67 |
| KEGG | 34517 | 74.05 |
| InterPro | 43884 | 94.14 |
| GO | 27155 | 58.25 |
| Pfam | 34383 | 73.76 |
| eggNOG | 42434 | 91.03 |
| Annotated | 45431 | 97.46 |
| Unannotated | 1185 | 2.54 |

**Table S13. The COG distribution of functional annotation of genes in Newhall navel orange genome**

| COG category | Gene number | Percentage (%) |
|---|---|---|
| A: RNA processing and modification | 1489 | 2.41 |
| B: Chromatin structure and dynamics | 580 | 0.94 |
| C: Energy production and conversion | 1547 | 2.50 |
| D: Cell cycle control, cell division, chromosome partitioning | 835 | 1.35 |
| E: Amino acid transport and metabolism | 1400 | 2.27 |
| F: Nucleotide transport and metabolism | 579 | 0.94 |
| G: Carbohydrate transport and metabolism | 2506 | 4.06 |
| H: Coenzyme transport and metabolism | 716 | 1.16 |
| I: Lipid transport and metabolism | 1379 | 2.23 |
| J: Translation, ribosomal structure, and biogenesis | 1983 | 3.21 |
| K: Transcription | 3600 | 5.83 |
| L: Replication, recombination, and repair | 1559 | 2.52 |
| M: Cell wall/membrane/envelop biogenesis | 838 | 1.36 |

| | | |
|---|---|---|
| N: Cell motility | 316 | 0.51 |
| O: Post-translational modification, protein turnover, and chaperones | 3388 | 5.48 |
| P: Inorganic ion transport and metabolism | 1357 | 2.20 |
| Q: Secondary metabolites biosynthesis, transport, and catabolism | 1848 | 2.99 |
| R: General function prediction only | 346 | 0.56 |
| S: Functional unknown | 23757 | 38.46 |
| T: Signal transduction mechanisms | 3905 | 6.32 |
| U: Intracellular trafficking, secretion, and vesicular transport | 1905 | 3.08 |
| NO COGs | 2249 | 3.64 |
| V: Defense mechanisms | 941 | 1.52 |
| W: Extracellular structures | 874 | 1.41 |
| X: Mobilome: prophages, transposons | 830 | 1.34 |
| Y: Nuclear structure | 267 | 0.43 |
| Z: Cytoskeleton | 783 | 1.27 |

**Table S14. The statistics of non-coding RNA annotation in Newhall navel orange genome**

| Type | | Copy number | Average length(bp) | Total length(bp) | Percentage of genome (%) |
|---|---|---|---|---|---|
| miRNA | | 931 | 109.73 | 102,162 | 0.014908 |
| tRNA | | 1,200 | 75.39 | 90,469 | 0.013202 |
| rRNA | rRNA | 6,765 | 332.31 | 2,248,109 | 0.33 |
| | 18S | 916 | 1,581.45 | 1,448,611 | 0.21 |
| | 28S | 3,153 | 142.14 | 448,180 | 0.065402 |
| | 5.8S | 803 | 161.31 | 129,531 | 0.018902 |
| | 5S | 1,893 | 117.16 | 221,787 | 0.032365 |
| snRNA | snRNA | 2,325 | 109.19 | 253,863 | 0.037045 |
| | CD-box | 2,076 | 105.77 | 219,575 | 0.032042 |
| | HACA-box | 78 | 131.95 | 10,292 | 0.001502 |
| | splicing | 168 | 140.27 | 23,566 | 0.003439 |
| | scaRNA | 3 | 143.33 | 430 | 0.000063 |
| | Unknown | 0 | 0 | 0 | 0 |

**Table S15. The species information of 26 accessions of citrus and relatives**

| Common name | Species name | Species group | Abb. | Genome status | HLB response based on Alves et al. 2021 | Accession number of genomic data | Coverage | Citations |
|---|---|---|---|---|---|---|---|---|
| Chinese box-orange | *Atlantia buxifolia* | Atlantia | HKC | Assembled in previous study | susceptible | PRJNA327148 | 401X | Wang et al. 2017 |
| Australian finger lime admixture | *Microcitrus australasica (F Muell.)* | Australian.limes | AFL | not assembled | resistant | PRJNA414519 | 69X | Wu et al. 2018 |
| Australian round lime | *Microcitrus australis Swingle* | Australian.limes | ARR | not assembled | resistant | PRJNA414519 | 25X | Wu et al. 2018 |
| Eremorange | *Eremocitrus glauca x C. sinensis* | Australian.limes | ADL | not assembled | resistant | PRJNA414519 | 59X | Wu et al. 2018 |
| Humpang citron | *C. medica* | Citron | HUM | not assembled | Tolerant | PRJNA414519 | 69X | Wu et al. 2018 |
| Mac Veu Mountain citron | *C. medica* | Citron | VEU | not assembled | Tolerant | PRJNA414519 | 63X | Wu et al. 2018 |
| Citron | *C. medica* | Citron | XZ | Assembled in previous study | Tolerant | PRJNA320023 | 344X | Wang et al. 2017 |
| Kumquat | *Fortunella hindsii* | Kumquat | FOR | Assembled in previous study | Tolerant | PRJNA487160 | 137X | Zhu et al. 2019 |
| Red rough lemon | *Citrus jambhiri Lush.* | Lemons | RRL | not assembled | Tolerant | PRJNA414519 | 68X | Wu et al. 2018 |
| Eureka lemon | *C. x limon L. (Burm. F.)* | Lemons | LIM | not assembled | Tolerant | PRJNA414519 | 114X | Wu et al. 2018 |
| Rangpur lime | *C. x limonia (Osbeck)* | Lime | LMA | not assembled | susceptible | PRJNA414519 | 59X | Wu et al. 2018 |
| Willowleaf mandarin | *C. × deliciosa* | Mandarins | WLM | Not assembled | susceptible | SRX372685 | 125X | Wu et al. 2014 |
| Dancy mandarin | *Citrus reticulata* | Mandarins | DNC | Not assembled | susceptible | PRJNA414519 | 60X | Wu et al. 2018 |
| Satsuma mandarin | *C. x unshiu (Marc)* | Mandarins | UNS | Not assembled | susceptible | PRJNA414519 | 63X | Wu et al. 2018 |
| Sunki mandarin | *C. x sunki (hayata)* | Mandarins | SNK | Not assembled | susceptible | PRJNA414519 | 63X | Wu et al. 2018 |
| Sun Chu Sha Kat | *Citrus reticulata* | Mandarins | SCM | Not assembled | susceptible | PRJNA414519 | 38X | Wu et al. 2018 |
| Tachibana mandarin | *Citrus reticulata* | Mandarins | TBM | Not assembled | susceptible | PRJNA414519 | 32X | Wu et al. 2018 |
| Cleopatra mandarin | *C. x reshni (Hort. ex Tanaka)* | Mandarins | CLP | Not assembled | susceptible | PRJNA414519 | 126X | Wu et al. 2018 |
| Clementine mandarin | *C. × clementina cv. Clemenules* | Mandarins | CLM | Assembled in previous study | susceptible | SRX371962 | 7X | Wu et al. 2014 |
| Mangshan mandarin | *Citrus reticulata* | Mandarins | CMS | Assembled in previous study | susceptible | PRJNA392792 | 35X | Wang et al. 2018 |
| Ichange papeda | *Citrus ichangensis* | Papeda | XJC | Assembled in previous study | Tolerant | PRJNA321657 | 164X | Wang et al. 2017 |

| | | | | | | | | |
|---|---|---|---|---|---|---|---|---|
| Low acid pummelo | *Citrus grandis* | Pummelo | LAP | Not assembled | susceptible | PRJNA225967 | 20X | Wu et al. 2018 |
| Pummelo | *Citrus grandis* | Pummelo | HWB | Assembled in previous study | susceptible | PRJNA318855 | 57X | Wang et al. 2017 |
| Newhall sweet orange | *Citrus sinensis* | Sweet orange | CSN | Assembled in this study | susceptible | PRJNA810206 | 50X | This study |
| Valencia sweet orange | *Citrus sinensis* | Sweet orange | CSV | Assembled in previous study | susceptible | PRJNA86123 | 100X | Xu et al. 2013 |
| Trifoliate orange | *Poncirus trifoliata* | Trifoliate orange | PON | Assembled in previous study | Tolerant | PRJNA648176 | 91X | Peng et al. 2020 |

Note: Yellow highlight indicates assembled citrus genomes.

**Table S16. COG distribution of pan, core, soft core, dispensable, and private genes in citrus genomes**

| COG category | core gene | | soft core gene | | dispensable gene | | private gene | | pan genome |
|---|---|---|---|---|---|---|---|---|---|
| | Gene number | p value[a] | Gene number | p value | Gene number | p value | Gene number | p value | Gene number |
| A: RNA processing and modification | 497 | 1.49E-16 | 101 | 0.976571 | 272 | 0.9996611 | 481 | 0.999897 | 1351 |
| B: Chromatin structure and dynamics | 141 | 0.00134306 | 27 | 0.97677 | 95 | 0.7686883 | 159 | 0.88607 | 422 |
| C: Energy production and conversion | 374 | 0.05311679 | 124 | 0.237126 | 329 | 0.1314954 | 475 | 0.998554 | 1302 |
| D: Cell cycle control, cell division, chromosome partitioning | 236 | 2.70E-06 | 42 | 0.996908 | 127 | 0.999634 | 276 | 0.492524 | 681 |
| E: Amino acid transport and metabolism | 411 | 5.04E-05 | 97 | 0.976987 | 288 | 0.945229 | 507 | 0.87642 | 1303 |
| F: Nucleotide transport and metabolism | 130 | 0.009898899 | 27 | 0.960645 | 94 | 0.6608604 | 155 | 0.835706 | 406 |
| G: Carbohydrate transport and metabolism | 776 | 3.15E-17 | 193 | 0.723674 | 476 | 0.9991478 | 799 | 0.999999 | 2244 |
| H: Coenzyme transport and metabolism | 147 | 0.02190847 | 54 | 0.040265 | 88 | 0.9981131 | 186 | 0.729146 | 475 |
| I: Lipid transport and metabolism | 393 | 1.27E-06 | 105 | 0.592825 | 252 | 0.9921721 | 447 | 0.987297 | 1197 |
| J: Translation, ribosomal structure and biogenesis | 511 | 9.29E-05 | 153 | 0.349176 | 362 | 0.9814447 | 633 | 0.974035 | 1659 |
| K: Transcription | 1238 | 3.17E-89 | 328 | 5.96E-07 | 491 | 1 | 777 | 1 | 2834 |
| L: Replication, recombination and repair | 430 | 0.09325893 | 118 | 0.956659 | 423 | 0.00023946 | 552 | 0.999686 | 1523 |
| M: Cell wall/membrane/envelop biogenesis | 194 | 4.57E-05 | 39 | 0.967741 | 111 | 0.994404 | 223 | 0.71747 | 567 |
| N: Cell motility | 2 | 0.6743357 | 0 | 1 | 0 | 1 | 6 | 0.052476 | 8 |
| O: Post-translational modification, protein turnover, and chaperones | 1107 | 2.78E-20 | 337 | 0.003467 | 755 | 0.9007754 | 1082 | 1 | 3281 |
| P: Inorganic ion transport and metabolism | 338 | 1.21E-07 | 108 | 0.017972 | 193 | 0.9996686 | 351 | 0.999466 | 990 |
| Q: Secondary metabolites biosynthesis, transport, and catabolism | 437 | 0.8976643 | 222 | 1.55E-08 | 438 | 0.06616386 | 622 | 0.999887 | 1719 |
| S: Functional unknown | 4712 | 5.20E-136 | 1968 | 6.42E-145 | 3502 | 1.84E-11 | 3267 | 1 | 13449 |
| T: Signal transduction mechanisms | 1098 | 6.89E-13 | 308 | 0.471187 | 869 | 0.02459055 | 1156 | 1 | 3431 |
| U: Intracellular trafficking, secretion, and vesicular transport | 509 | 4.16E-15 | 101 | 0.9936 | 291 | 0.9987944 | 511 | 0.999571 | 1412 |
| NO COGs | 15 | 1 | 161 | 1 | 2970 | 3.78E-09 | 8286 | 0 | 11432 |
| V: Defense mechanisms | 114 | 0.000615924 | 48 | 0.000426 | 80 | 0.4176458 | 84 | 1 | 326 |
| W: Extracellular structures | 35 | 4.06E-05 | 8 | 0.299501 | 11 | 0.9696151 | 17 | 0.998852 | 71 |
| Y: Nuclear structure | 39 | 0.002651072 | 4 | 0.97784 | 10 | 0.9998143 | 44 | 0.186928 | 97 |
| Z: Cytoskeleton | 195 | 5.63E-09 | 33 | 0.981571 | 77 | 0.9999996 | 203 | 0.601384 | 508 |

[a]Fisher's exact test.

**Table S17. The statistics of structural variants (SVs) for 25 citrus genomes**

| Species | Deletions | Insertions | SNP |
|---|---|---|---|
| ADL | 4,098 | 3,654 | 6,256,077 |
| AFL | 4,851 | 4,517 | 6,089,282 |
| ARR | 5,958 | 5,555 | 5,441,333 |
| CLM | 4,457 | 4,557 | 3,309,221 |
| CLP | 6,248 | 5,582 | 3,675,605 |
| DNC | 5,856 | 5,254 | 3,890,748 |
| FOR | 4,969 | 5,231 | 5,241,039 |
| HKC | 404 | 402 | 4,994,012 |
| HUM | 7,864 | 7,117 | 5,485,865 |
| HWB | 4,279 | 4,678 | 3,480,724 |
| LAP | 12,107 | 10,538 | 3,715,682 |
| LIM | 5,381 | 4,915 | 6,136,827 |
| LMA | 4,630 | 4,206 | 6,256,077 |
| CMS | 6,126 | 6,035 | 3,610,472 |
| RRL | 4,649 | 4,240 | 6,046,652 |
| SCM | 5,865 | 5,300 | 3,861,637 |
| SNK | 6,177 | 5,552 | 3,782,008 |
| TBM | 15,881 | 13,961 | 3,719,227 |
| CSN | 4,464 | 4,895 | 3,944,830 |
| UNS | 5,658 | 5,009 | 3,897,032 |
| VEU | 7,965 | 7,221 | 5,417,629 |
| WLM | 5,653 | 5,098 | 3,877,263 |
| XJC | 5,837 | 6,970 | 4,244,201 |
| XZ | 4,344 | 5,050 | 5,140,422 |
| PON | 2,475 | 2,436 | 5,931,022 |

**Table S18. COG distribution of unique genes for HLB-susceptible and -tolerant citrus genomes**

| COG category | HLB tolerant genotype | | HLB susceptible genotype | | pan genome |
|---|---|---|---|---|---|
| | Gene number | p value[a] | Gene number | p value | Gene number |
| A: RNA processing and modification | 154 | 0.9777836 | 398 | 0.9995501 | 1351 |
| B: Chromatin structure and dynamics | 49 | 0.3067596 | 140 | 0.9596017 | 422 |
| C: Energy production and conversion | 199 | 0.9998033 | 358 | 0.1903944 | 1302 |
| D: Cell cycle control, cell division, chromosome partitioning | 80 | 0.2777913 | 225 | 0.9815981 | 681 |
| E: Amino acid transport and metabolism | 206 | 0.9950219 | 374 | 0.07883045 | 1303 |
| F: Nucleotide transport and metabolism | 74 | 0.9968285 | 105 | 0.01884531 | 406 |
| G: Carbohydrate transport and metabolism | 273 | 0.9993458 | 648 | 0.9992679 | 2244 |
| H: Coenzyme transport and metabolism | 87 | 0.9956376 | 126 | 0.01045465 | 475 |
| I: Lipid transport and metabolism | 153 | 0.9228342 | 360 | 0.9532799 | 1197 |
| J: Translation, ribosomal structure and biogenesis | 242 | 0.9883502 | 488 | 0.4261893 | 1659 |
| K: Transcription | 235 | 1 | 650 | 1 | 2834 |
| L: Replication, recombination and repair | 180 | 0.8519681 | 468 | 0.998755 | 1523 |
| M: Cell wall/membrane/envelop biogenesis | 93 | 0.9966397 | 152 | 0.09811826 | 567 |
| N: Cell motility | 4 | 0.7808764 | 2 | 0.01854325 | 8 |
| O: Post-translational modification, protein turnover, and chaperones | 346 | 1 | 911 | 1 | 3281 |
| P: Inorganic ion transport and metabolism | 114 | 0.9917783 | 282 | 0.9968407 | 990 |
| Q: Secondary metabolites biosynthesis, transport, and catabolism | 259 | 0.9999758 | 473 | 0.2229415 | 1719 |
| S: Functional unknown | 1249 | 1 | 2879 | 1 | 13449 |
| T: Signal transduction mechanisms | 373 | 0.9999307 | 996 | 1 | 3431 |
| U: Intracellular trafficking, secretion, and vesicular transport | 193 | 0.9995976 | 394 | 0.7982337 | 1412 |
| NO COGs | 2912 | 0 | 6117 | 6.68E-284 | 11432 |
| V: Defense mechanisms | 41 | 0.9999998 | 64 | 0.8470419 | 326 |
| W: Extracellular structures | 4 | 0.9841154 | 15 | 0.9943368 | 71 |
| Y: Nuclear structure | 10 | 0.1614453 | 36 | 0.9072949 | 97 |
| Z: Cytoskeleton | 60 | 0.5991702 | 160 | 0.9616464 | 508 |

[a]Fisher's exact test.

**Table S19. Group specific SNP and Indel analyses of genes involved in plant immunity for 26 citrus accessions in three groups (HLB-resistant, -tolerant, and -susceptible)**

| Group | Genes | Annotation | Detailed annotation | InDels number | Mutations on CDS | Mutations on protein |
|---|---|---|---|---|---|---|
| Resistant | Cs_ont_1g002940.1 | NBS-LRR | XP_006427948.2, disease resistance protein | 3 | c.1623_1625delcct, c.1970t > atgactacaa | p.Leu542del, p.Thr656_Yyr657insAspAspTyr,p.Val657Asn |
| Resistant | Cs_ont_1g011240.1 | NBS-LRR | KAH9715781.1, disease resistance protein | 1 | c.1732insg | p.Gln578Alafs*619 |
| Resistant | Cs_ont_1g015380.1 | NBS-LRR | XP_015386355.1, disease resistance protein | 6 | c.369detg, c.654insctt, c.696_732deltgtgttggaaatatttccacatagattctttgaatct, c.1172_1173delcg, c.1129dela, c.1624_1648instggtgggagcatggtgggagcagtt | p.Glu125del,p.Asp130* |
| Resistant | Cs_ont_1g015390.1 | NBS-LRR | KAH9714655.1, disease resistance protein | 4 | c.2178_2183delaaggac | p.Met1fs*7 |
| Resistant | Cs_ont_1g018870.1 | NBS-LRR | XP_006465867.1, disease resistance protein | 6 | c.177_178delcgtgaag, c.714_761deltagggatgagatcagggttgctagggatgagatcagtgttgctaggga, c.933delC, c.998_999ga>ag, c.2863insaaa | p.Asp59fs*67 |
| Resistant | Cs_ont_1g020300.1 | NBS-LRR | XP_006466045.1, disease resistance protein | 3 | c.718insaatgtc, c.722_723aa>tg, c.1462insgaa | p.Asp239_Asp240insAsnVal |
| Resistant | Cs_ont_2g015810.1 | NBS-LRR | KAH9745578.1,Disease resistance protein | 3 | c.1098delgga, c.1103t>a, c.1105_1106aa>gt | p.Val366_Leu367insGlu, p.Val368Glu, p.Asn3369Val |
| Resistant | Cs_ont_2g031880.1 | NBS-LRR | XP_024950205.1, disease resistance protein RPP13-like | 1 | c.1555_1563deltgcaaggat, | p.Cys519_Asp521del |
| Resistant | Cs_ont_2g031900.1 | NBS-LRR | XP_006470643.1, disease resistance protein RPP13-like | 2 | c.1940_1944tttct>a, c.1946a>g | p.Phe647* |
| Resistant | Cs_ont_3g021990.1 | NBS-LRR | KAH9753096.1, Disease resistance protein | 2 | c.1962_1964gaa>att | p.Lys565Leu |
| Resistant | Cs_ont_3g029080.1 | NBS-LRR | XP_006471150.1, disease resistance protein | 1 | c.3434t>c, | p.Leu1145Pro,p.His1145_*1146delinsLeuAsnValPheProSerThrLeu |
| Resistant | Cs_ont_3g029450.1 | NBS-LRR | XP_006471132.1, disease resistance protein | 2 | c.387_423delctttaatacgacttcctacactcctgctccagagtgg, c.1136insgtt | p.Asn129fs*155 |
| Resistant | Cs_ont_4g014690.1 | NBS-LRR | XP_024034355.1, disease resistance protein | 1 | c.1809instgaatc | p.Asp603_Glu604insGluSer |

| | | | | | | |
|---|---|---|---|---|---|---|
| Resistant | Cs_ont_4g015140.1 | NBS-LRR | XP_024949266.1, disease resistance protein | 10 | c.758_759deltg, c.764c>t, c.1107_1109ccg>ggc, c.1517delc, c.1657_1658delag, c.1737_1741delaaggc, c.2182_2184caa>atg, c.2187insg | p.Met253fs*257 |
| Resistant | Cs_ont_5g019650.1 | NBS-LRR | XP_006478606.1, disease resistance protein | 1 | c.2015insa | p.Ser673fs*738 |
| Resistant | Cs_ont_5g020660.1 | NBS-LRR | XP_006442968.1, disease resistance protein | 1 | c.448insC | p.Glu150_Arg151delinsArgThrAla* |
| Resistant | Cs_ont_5g021910.1 | NBS-LRR | KAH9782658.1,putative disease resistance protein | 3 | c.729_731aga>cag,1080insC | p.Glu244Arg,p.Phe361fs*367 |
| Resistant | Cs_ont_7g017950.1 | NBS-LRR | XP_024956129.1,putative disease resistance protein | 2 | c.621a>t,c.626_646delaaaagcaagctaatgaaaatg | p.Lys207Asn,p.Lys210_Glu216del |
| Resistant | Cs_ont_8g005070.1 | NBS-LRR | KAH9706177.1,Disease resistance protein | 2 | c.340delt,c.404insG | p.Phe115fs*123 |
| Resistant | Cs_ont_8g019180.1 | NBS-LRR | KAH9657265.1,Disease resistance protein | 1 | c.3280inscagagag | p.Ile1094fs*1127 |
| Resistant | Cs_ont_8g019690.1 | NBS-LRR | XP_024957917.1,disease resistance protein | 1 | c.3207insgg | p.Asp1070fs*1094 |
| Resistant | Cs_ont_8g019780.1 | NBS-LRR | KAH9696680.1,Disease resistance protein | 12 | c.1_11delatggaaatttc,c.14c>g,c.16_17ct>tc,c.28_40cttttagaaccga>aag,c.164_165delag,c.825insta,c.825_826cg>ta,c.828_831ttct>gcga,c.833_836tcga>gtcg, | p.Met1_Ile14delinsMetIlePheArgThrLysVal,p.Arg56fs*59 |
| Resistant | Cs_ont_8g019940.1 | NBS-LRR | KAH9696663.1,Disease resistance protein | 4 | c.10a>t,c.14t>a,c.17g>a,c.26_44aagactgtttaatagactg>t,1019_1020delgg, | p.Ile4_Cys4del,p.Asp14_Cys15del,p.Arg340fs*344 |
| Resistant | Cs_ont_8g019960.1 | NBS-LRR | KAH9696658.1,Disease resistance protein | 5 | c.1900_1904cgaaa>gatgg,c.2261_2263deltat,c.2945dela | p.Arg634Asp,p.Asn635Gly,p.Leu755del,p.Asp983fsAsn998 |
| Resistant | Cs_ont_8g019970.1 | NBS-LRR | KAH9696656.1,Disease resistance protein | 2 | c.1080_1084delaaaag | p.Lys361fs*365 |
| Resistant | Cs_ont_8g019980.1 | NBS-LRR | KAH9696656.1,Disease resistance protein | 2 | c.363_367deltaaat,c.466_477delgaagatgccaat | p.Lys122* |
| Resistant | Cs_ont_5g031840.1 | receptor-like kinase | XP_006493305.1,receptor-like protein kinase | 1 | c.2529_2532aaac>tgaa | p.Asn846Glu |
| Resistant | Cs_ont_6g016380.1 | receptor-like kinase | KAH9672564.1,Receptor-like serine/threonine-protein kinase | 1 | c.1715inst | p.Gly572fs*575 |
| Resistant | Cs_ont_5g031860.1 | receptor-like kinase | KAH9779927.1,Receptor-like protein kinase | 1 | c.625_627delcgg | p.Arg209del |
| Resistant | Cs_ont_8g006180.1 | SOD | XP_006488000.1,superoxide dismutase [Cu-Zn], chloroplastic | 1 | c.115_117deltct | p.Ser43del |

| | | | | | | |
|---|---|---|---|---|---|---|
| Tolerant | Cs_ont_3g016780.1 | NBS-LRR | XP_006471944.1,probable disease resistance protein | 1 | c.2372insaacctt | p.Asn911delinsLysProTyr |
| Tolerant | Cs_ont_8g019940.1 | NBS-LRR | KAH9696663.1,Disease resistance protein | 1 | c.1014insgcg | p.Leu338_Lys339insArg |
| Tolerant | Cs_ont_2g031900.1 | NBS-LRR | XP_006470643.1,disease resistance protein | 2 | c.397_398at>acttccaatgagaagctcgaaatggattgaaggaagcac | p.Ile133Thr,p.Gly135fs*143 |
| Tolerant | Cs_ont_5g041000.1 | SOD | XP_006477334.1,copper chaperone for superoxide dismutase, chloroplastic/cytosolic | 1 | c.73inscсttcttct | p.Phe24_Ser25insProSerSer |
| Susceptible | Cs_ont_4g015140.1 | NBS-LRR | XP_024949266.1,probable disease resistance protein | 1 | c.1040insgg | p.Lys347fs*357 |
| Susceptible | Cs_ont_1g017970.1 | receptor-like kinase | XP_006465723.3,putative leucine-rich repeat receptor-like protein kinase | 1 | c.239_240delcg | p.Ser80fs*95 |

Note: c. cDNA reference sequence; c.123A>T: compared to the reference sequence, A123 was replaced by T; c.2052delA: compared with the reference, A2052 was deleted; c.5756_5757insAGG: compared with the reference, AGG was inserted between 5756 and 5757; p.: reference sequence of protein; p.Ala3_Ser5del: Ala3 to Ser5 were deleted; p.Lys2_Gly3insGlnSerLys: GlnSerLys was inserted between Lys2 and Gly3; p.Arg97ProfsTer99: Arg97 was the mutated into Pro. In addition, frame shift happened that led to stop codon at 99; p.Asn846Glu: Asn846 was replaced by Glu.

**Table S20. The species information of citrus and relatives accessions for GWAS study**

| Group | HLB | Common name | Species name | Data accession | Genome status | Sequencing technology | Reference |
|---|---|---|---|---|---|---|---|
| Australian lime | Resistant/ Tolerant | Australian round lime 1 | *Microcitrus australis Swingle* | PRJNA414519 | raw reads | Illumina | Wu et al., 2018 |
| | | Australian round lime 2 | *Microcitrus australis Swingle* | PRJNA414519 | raw reads | Illumina | Wu et al., 2018 |
| | | Microcitrus Swingle | *Microcitrus sp.* | http://citgvd.cric.cn/home/index | raw reads | Illumina | Li et al., 2020 |
| | | Australian finger lime | *Microcitrus australasica (F Muell.)* | PRJNA414519 | raw reads | Illumina | Wu et al., 2018 |
| | | Australian finger lime admicture | *Microcitrus australasica (F Muell.)* | PRJNA414519 | raw reads | Illumina | Wu et al., 2018 |
| | | Eremocitrus Swingle | *Eremocitrus sp.* | http://citgvd.cric.cn/home/index | raw reads | Illumina | Li et al., 2020 |
| | | Australian desert lime | *Eremocitrus glauca* | PRJNA414519 | raw reads | Illumina | Wu et al., 2018 |
| | | Eremorange | *Eremocitrus glauca x C. sinensis* | PRJNA414519 | raw reads | Illumina | Wu et al., 2018 |
| Citron | Resistant/ Tolerant | Citron | *Citrus medica* | PRJNA320023 | Draft assemblied genome | Illumina | Wang et al., 2017 |
| | | wild citron XZ | *C. medica* | PRJNA320023 | raw reads | Illumina | Wang et al., 2017 |
| | | wild citron XZ1 | *C. medica* | PRJNA320023 | raw reads | Illumina | Wang et al., 2017 |
| | | wild citron JY | *C. medica* | PRJNA320023 | raw reads | Illumina | Wang et al., 2017 |
| | | wild citron JY4 | *C. medica* | PRJNA320023 | raw reads | Illumina | Wang et al., 2017 |
| | | wild citron JY5 | *C. medica* | PRJNA320023 | raw reads | Illumina | Wang et al., 2017 |
| | | wild citron JY8 | *C. medica* | PRJNA320023 | raw reads | Illumina | Wang et al., 2017 |
| | | wild citron JY15 | *C. medica* | PRJNA320023 | raw reads | Illumina | Wang et al., 2017 |
| | | wild citron JY28 | *C. medica* | PRJNA320023 | raw reads | Illumina | Wang et al., 2017 |
| | | wild citron FS | *C. medica* | PRJNA320023 | raw reads | Illumina | Wang et al., 2017 |
| | | Corsican citron | *C. medica L.* | PRJNA414519 | raw reads | Illumina | Wu et al., 2018 |
| | | Mac Veu citron | *C. medica L.* | PRJNA414519 | raw reads | Illumina | Wu et al., 2018 |
| | | Buddha´s hand citron | *C. medica L.* | PRJNA414519 | raw reads | Illumina | Wu et al., 2018 |
| | | Humpang citron | *C. medica L.* | PRJNA414519 | raw reads | Illumina | Wu et al., 2018 |

| | | | | | | | |
|---|---|---|---|---|---|---|---|
| | | Foshou | *Citras medica L.* | http://citgvd.cric.cn/home/index | raw reads | Illumina | Li et al., 2020 |
| | | Mexico Citron | *Citrus medica L* | http://citgvd.cric.cn/home/index | raw reads | Illumina | Li et al., 2020 |
| | | Arizona 861 | *Citrus medica L* | http://citgvd.cric.cn/home/index | raw reads | Illumina | Li et al., 2020 |
| | | Guangxituningmeng | *Citrus sp.* | http://citgvd.cric.cn/home/index | raw reads | Illumina | Li et al., 2020 |
| | | Leimeng | *Citrus sp.* | http://citgvd.cric.cn/home/index | raw reads | Illumina | Li et al., 2020 |
| Lemon | Resistant/ Tolerant | Eureka lemon | *C. x limon L. (Burm. F.)* | PRJNA414519 | raw reads | Illumina | Wu et al., 2018 |
| | | Lemon | *C.* × *limon* | PRJNA320023 | raw reads | Illumina | Wang et al., 2017 |
| | | Lemon | *C.* × *limon* | PRJNA320023 | raw reads | Illumina | Wang et al., 2017 |
| | | Lemon | *C.* × *limon* | PRJNA320023 | raw reads | Illumina | Wang et al., 2017 |
| | | Lemon | *C.* × *limon* | PRJNA320023 | raw reads | Illumina | Wang et al., 2017 |
| | | Lemon | *C.* × *limon* | PRJNA320023 | raw reads | Illumina | Wang et al., 2017 |
| | | Lemon | *C.* × *limon* | PRJNA320023 | raw reads | Illumina | Wang et al., 2017 |
| | | Lemon | *C.* × *limon* | PRJNA320023 | raw reads | Illumina | Wang et al., 2017 |
| | | improved_meyer_lemon | *Citrus x meyeri 'Improved'* | | raw reads | BGISEQ500 | In this study |
| | | volckameriana | *C.* × *limon* | PRJNA698060 | raw reads | BGISEQ500 | In this study |
| | | volckameriana | *C.* × *limon* | PRJNA698060 | raw reads | BGISEQ500 | In this study |
| | | volckameriana | *C.* × *limon* | PRJNA698060 | raw reads | BGISEQ500 | In this study |
| | | volckameriana | *C.* × *limon* | PRJNA698060 | raw reads | BGISEQ500 | In this study |
| | | volckameriana | *C.* × *limon* | PRJNA698060 | raw reads | BGISEQ500 | In this study |
| | | volckameriana | *C.* × *limon* | PRJNA698060 | raw reads | BGISEQ500 | In this study |
| | | volckameriana | *C.* × *limon* | PRJNA698060 | raw reads | BGISEQ500 | In this study |
| | | volckameriana | *C.* × *limon* | PRJNA698060 | raw reads | BGISEQ500 | In this study |

| | | | | | | | |
|---|---|---|---|---|---|---|---|
| Ponderosa lemon | | ponderosa_lemon | *C. x pyriformis* | PRJNA698060 | raw reads | BGISEQ500 | In this study |
| Rough lemon | | rough_lemon | *Citrus jambhiri Lush.* | PRJNA698060 | raw reads | BGISEQ500 | In this study |
| | | rough_lemon | *Citrus jambhiri Lush.* | PRJNA698060 | raw reads | BGISEQ500 | In this study |
| | | rough_lemon | *Citrus jambhiri Lush.* | PRJNA698060 | raw reads | BGISEQ500 | In this study |
| | | Red rough lemon | *C. x jambhiri (Lush)* | PRJNA414519 | raw reads | Illumina | Wu et al., 2018 |
| Kaffir lime | susceptible | Kaffir lime | *C. hystrix* | PRJNA698060 | raw reads | Illumina | In this study |
| Rangpur lime | | Honglimeng | *Citrus limonia* | http://citgvd.cric.cn/home/index | raw reads | Illumina | Li et al., 2020 |
| | | Rangpur lime | *C. x limonia (Osbeck)* | PRJNA414519 | raw reads | Illumina | Wu et al., 2018 |
| Mexican lime | | Mexican lime | *C. x aurantifolia (Christm.) Swingle* | PRJNA414519 | raw reads | Illumina | Wu et al., 2018 |
| | | mexican_lime | *Citrus x aurantifolia* | PRJNA698060 | raw reads | BGISEQ500 | In this study |
| Sour orange | | Jiangjinsuancheng | *Citrus aurantium L.* | http://citgvd.cric.cn/home/index | raw reads | Illumina | Li et al., 2020 |
| | | Zhuluan | *Citrus aurantium L.* | http://citgvd.cric.cn/home/index | raw reads | Illumina | Li et al., 2020 |
| | | Daidai | *Citrus aurantium L.* | http://citgvd.cric.cn/home/index | raw reads | Illumina | Li et al., 2020 |
| | | Goutoucheng | *Citrus aurantium L.* | http://citgvd.cric.cn/home/index | raw reads | Illumina | Li et al., 2020 |
| | | sour orange | *C. x aurantium, L.* | PRJNA321100 | raw reads | Illumina | Wang et al., 2017 |
| | | sour orange | *C. x aurantium, L.* | PRJNA321100 | raw reads | Illumina | Wang et al., 2017 |
| | | sour orange | *C. x aurantium, L.* | PRJNA321100 | raw reads | Illumina | Wang et al., 2017 |
| | | sour orange | *C. x aurantium, L.* | PRJNA321100 | raw reads | Illumina | Wang et al., 2017 |
| | | sour orange | *C. x aurantium, L.* | PRJNA321100 | raw reads | Illumina | Wang et al., 2017 |
| | | sour orange | *C. x aurantium, L.* | PRJNA321100 | raw reads | Illumina | Wang et al., 2017 |
| | | Seville sour orange | *C. x aurantium, L.* | SRX372786 | raw reads | Illumina | Wu et al., 2014 |
| Sweet orange | | Sweet Orange | *C. sinensis* | PRJNA86123 | Draft assemblied genome | Illumina | Xu et al., 2013 |

| | | | | | | | |
|---|---|---|---|---|---|---|---|
| | | sweet-orange variety 'Ridge Pineapple' | *C. sinensis* | SRX372703 | Draft assemblied genome | Sanger/454 | Wu et al., 2014 |
| | | Sweet Orange | *C. sinensis* | PRJNA321100 | raw reads | Illumina | Wang et al., 2017 |
| | | Sweet Orange | *C. sinensis* | PRJNA321100 | raw reads | Illumina | Wang et al., 2017 |
| | | Sweet Orange | *C. sinensis* | PRJNA321100 | raw reads | Illumina | Wang et al., 2017 |
| | | Sweet Orange | *C. sinensis* | PRJNA321100 | raw reads | Illumina | Wang et al., 2017 |
| | | Sweet Orange | *C. sinensis* | PRJNA321100 | raw reads | Illumina | Wang et al., 2017 |
| | | Sweet Orange | *C. sinensis* | PRJNA321100 | raw reads | Illumina | Wang et al., 2017 |
| | | Sweet Orange | *C. sinensis* | PRJNA321100 | raw reads | Illumina | Wang et al., 2017 |
| | | Sweet Orange | *C. sinensis* | PRJNA321100 | raw reads | Illumina | Wang et al., 2017 |
| | | Sweet Orange | *C. sinensis* | PRJNA321100 | raw reads | Illumina | Wang et al., 2017 |
| | | Sweet Orange | *C. sinensis* | PRJNA321100 | raw reads | Illumina | Wang et al., 2017 |
| | | Sweet Orange | *C. sinensis* | PRJNA321100 | raw reads | Illumina | Wang et al., 2017 |
| | | Sweet Orange | *C. sinensis* | PRJNA321100 | raw reads | Illumina | Wang et al., 2017 |
| | | Sweet Orange | *C. sinensis* | PRJNA321100 | raw reads | Illumina | Wang et al., 2017 |
| | | Sweet Orange | *C. sinensis* | PRJNA321100 | raw reads | Illumina | Wang et al., 2017 |
| | | Sweet Orange | *C. sinensis* | PRJNA321100 | raw reads | Illumina | Wang et al., 2017 |
| | | Sweet Orange | *C. sinensis* | PRJNA321100 | raw reads | Illumina | Wang et al., 2017 |
| | | Sweet Orange | *C. sinensis* | PRJNA321100 | raw reads | Illumina | Wang et al., 2017 |
| | | Sweet Orange | *C. sinensis* | PRJNA321100 | raw reads | Illumina | Wang et al., 2017 |
| | | Sweet Orange | *C. sinensis* | PRJNA321100 | raw reads | Illumina | Wang et al., 2017 |
| | | Sweet Orange | *C. sinensis* | PRJNA321100 | raw reads | Illumina | Wang et al., 2017 |
| | | Sweet Orange | *C. sinensis* | PRJNA321100 | raw reads | Illumina | Wang et al., 2017 |

| | | | | | | | |
|---|---|---|---|---|---|---|---|
| | | Sweet Orange | *C. sinensis* | PRJNA321100 | raw reads | Illumina | Wang et al., 2017 |
| | | Sweet Orange | *C. sinensis* | PRJNA321100 | raw reads | Illumina | Wang et al., 2017 |
| | | Sweet Orange | *C. sinensis* | PRJNA321100 | raw reads | Illumina | Wang et al., 2017 |
| | | Sweet Orange | *C. sinensis* | PRJNA321100 | raw reads | Illumina | Wang et al., 2017 |
| | | Sweet Orange | *C. sinensis* | PRJNA321100 | raw reads | Illumina | Wang et al., 2017 |
| | | Sweet Orange | *C. sinensis* | PRJNA321100 | raw reads | Illumina | Wang et al., 2017 |
| | | Sweet Orange | *C. sinensis* | PRJNA321100 | raw reads | Illumina | Wang et al., 2017 |
| | | Sweet Orange | *C. sinensis* | PRJNA321100 | raw reads | Illumina | Wang et al., 2017 |
| | | Sweet Orange | *C. sinensis* | PRJNA321100 | raw reads | Illumina | Wang et al., 2017 |
| | | Sweet Orange | *C. sinensis* | PRJNA321100 | raw reads | Illumina | Wang et al., 2017 |
| | | Sweet Orange | *C. sinensis* | PRJNA321100 | raw reads | Illumina | Wang et al., 2017 |
| | | Sweet Orange | *C. sinensis* | PRJNA321100 | raw reads | Illumina | Wang et al., 2017 |
| | | Sweet Orange | *C. sinensis* | PRJNA321100 | raw reads | Illumina | Wang et al., 2017 |
| | | Sweet Orange | *C. sinensis* | PRJNA321100 | raw reads | Illumina | Wang et al., 2017 |
| | | Sweet Orange | *C. sinensis* | PRJNA321100 | raw reads | Illumina | Wang et al., 2017 |
| | | Sweet Orange | *C. sinensis* | PRJNA321100 | raw reads | Illumina | Wang et al., 2017 |
| | | Sweet Orange | *C. sinensis* | PRJNA321100 | raw reads | Illumina | Wang et al., 2017 |
| | | Sweet Orange | *C. sinensis* | PRJNA321100 | raw reads | Illumina | Wang et al., 2017 |
| | | Sweet Orange | *C. sinensis* | PRJNA321100 | raw reads | Illumina | Wang et al., 2017 |
| | | Sweet Orange | *C. sinensis* | PRJNA321100 | raw reads | Illumina | Wang et al., 2017 |
| | | Sweet Orange | *C. sinensis* | PRJNA321100 | raw reads | Illumina | Wang et al., 2017 |
| | | Sweet Orange | *C. sinensis* | PRJNA321100 | raw reads | Illumina | Wang et al., 2017 |

| | | | | | | | |
|---|---|---|---|---|---|---|---|
| | | Sweet Orange | *C. sinensis* | PRJNA321100 | raw reads | Illumina | Wang et al., 2017 |
| | | Sweet Orange | *C. sinensis* | PRJNA321100 | raw reads | Illumina | Wang et al., 2017 |
| | | Sweet Orange | *C. sinensis* | PRJNA321100 | raw reads | Illumina | Wang et al., 2017 |
| | | Sweet Orange | *C. sinensis* | PRJNA321100 | raw reads | Illumina | Wang et al., 2017 |
| | | Sweet Orange | *C. sinensis* | PRJNA321100 | raw reads | Illumina | Wang et al., 2017 |
| | | Sweet Orange | *C. sinensis* | PRJNA321100 | raw reads | Illumina | Wang et al., 2017 |
| | | Sweet Orange | *C. sinensis* | PRJNA321100 | raw reads | Illumina | Wang et al., 2017 |
| | | Sweet Orange | *C. sinensis* | PRJNA321100 | raw reads | Illumina | Wang et al., 2017 |
| | | Sweet Orange | *C. sinensis* | PRJNA321100 | raw reads | Illumina | Wang et al., 2017 |
| | | Sweet Orange | *C. sinensis* | PRJNA321100 | raw reads | Illumina | Wang et al., 2017 |
| | | Sweet Orange | *C. sinensis* | PRJNA321100 | raw reads | Illumina | Wang et al., 2017 |
| | | Sweet Orange | *C. sinensis* | PRJNA321100 | raw reads | Illumina | Wang et al., 2017 |
| | | Sweet Orange | *C. sinensis* | PRJNA321100 | raw reads | Illumina | Wang et al., 2017 |
| | | Sweet Orange | *C. sinensis* | PRJNA321100 | raw reads | Illumina | Wang et al., 2017 |
| | | Sweet Orange | *C. sinensis* | PRJNA321100 | raw reads | Illumina | Wang et al., 2017 |
| | | Sweet Orange | *C. sinensis* | PRJNA321100 | raw reads | Illumina | Wang et al., 2017 |
| | | Sweet Orange | *C. sinensis* | PRJNA321100 | raw reads | Illumina | Wang et al., 2017 |
| | | Sweet Orange | *C. sinensis* | PRJNA321100 | raw reads | Illumina | Wang et al., 2017 |
| | | Sweet Orange | *C. sinensis* | PRJNA321100 | raw reads | Illumina | Wang et al., 2017 |
| | | Sweet Orange | *C. sinensis* | PRJNA321100 | raw reads | Illumina | Wang et al., 2017 |
| | | Sweet Orange | *C. sinensis* | PRJNA321100 | raw reads | Illumina | Wang et al., 2017 |
| | | Sweet Orange | *C. sinensis* | PRJNA321100 | raw reads | Illumina | Wang et al., 2017 |

| | | | | | | | |
|---|---|---|---|---|---|---|---|
| | | Sweet Orange | *C. sinensis* | PRJNA321100 | raw reads | Illumina | Wang et al., 2017 |
| | | Sweet Orange | *C. sinensis* | PRJNA321100 | raw reads | Illumina | Wang et al., 2017 |
| | | Sweet Orange | *C. sinensis* | PRJNA321100 | raw reads | Illumina | Wang et al., 2017 |
| | | Sweet Orange | *C. sinensis* | PRJNA321100 | raw reads | Illumina | Wang et al., 2017 |
| | | Sweet Orange | *C. sinensis* | PRJNA321100 | raw reads | Illumina | Wang et al., 2017 |
| | | Sweet Orange | *C. sinensis* | PRJNA321100 | raw reads | Illumina | Wang et al., 2017 |
| | | Sweet Orange | *C. sinensis* | PRJNA321100 | raw reads | Illumina | Wang et al., 2017 |
| | | Sweet Orange | *C. sinensis* | PRJNA321100 | raw reads | Illumina | Wang et al., 2017 |
| | | Sweet Orange | *C. sinensis* | PRJNA321100 | raw reads | Illumina | Wang et al., 2017 |
| | | Sweet Orange | *C. sinensis* | PRJNA321100 | raw reads | Illumina | Wang et al., 2017 |
| | | Sweet Orange | *C. sinensis* | PRJNA321100 | raw reads | Illumina | Wang et al., 2017 |
| | | Sweet Orange | *C. sinensis* | PRJNA321100 | raw reads | Illumina | Wang et al., 2017 |
| | | Sweet Orange | *C. sinensis* | PRJNA321100 | raw reads | Illumina | Wang et al., 2017 |
| | | Sweet Orange | *C. sinensis* | PRJNA321100 | raw reads | Illumina | Wang et al., 2017 |
| | | Sweet Orange | *C. sinensis* | PRJNA321100 | raw reads | Illumina | Wang et al., 2017 |
| | | Sweet Orange | *C. sinensis* | PRJNA321100 | raw reads | Illumina | Wang et al., 2017 |
| | | Sweet Orange | *C. sinensis* | PRJNA321100 | raw reads | Illumina | Wang et al., 2017 |
| | | Sweet Orange | *C. sinensis* | PRJNA321100 | raw reads | Illumina | Wang et al., 2017 |
| | | Sweet Orange | *C. sinensis* | PRJNA321100 | raw reads | Illumina | Wang et al., 2017 |
| | | Sweet Orange | *C. sinensis* | PRJNA321100 | raw reads | Illumina | Wang et al., 2017 |
| | | Sweet Orange | *C. sinensis* | PRJNA321100 | raw reads | Illumina | Wang et al., 2017 |
| | | Sweet Orange | *C. sinensis* | PRJNA321100 | raw reads | Illumina | Wang et al., 2017 |

| | | | | | | | |
|---|---|---|---|---|---|---|---|
| | | Sweet Orange | *C. sinensis* | PRJNA321100 | raw reads | Illumina | Wang et al., 2017 |
| | | Sweet Orange | *C. sinensis* | PRJNA321100 | raw reads | Illumina | Wang et al., 2017 |
| | | Sweet Orange | *C. sinensis* | PRJNA321100 | raw reads | Illumina | Wang et al., 2017 |
| | | Sweet Orange | *C. sinensis* | PRJNA321100 | raw reads | Illumina | Wang et al., 2017 |
| | | Sweet Orange | *C. sinensis* | PRJNA321100 | raw reads | Illumina | Wang et al., 2017 |
| | | Sweet Orange | *C. sinensis* | PRJNA321100 | raw reads | Illumina | Wang et al., 2017 |
| | | Sweet Orange | *C. sinensis* | PRJNA321100 | raw reads | Illumina | Wang et al., 2017 |
| | | Sweet Orange | *C. sinensis* | PRJNA321100 | raw reads | Illumina | Wang et al., 2017 |
| | | Sweet Orange | *C. sinensis* | PRJNA321100 | raw reads | Illumina | Wang et al., 2017 |
| Tangor | | algerian_clementine | *Citrus nobilis* | PRJNA414519 | raw reads | Illumina | Wu et al., 2018 |
| Grapefruit | | Grapefruit | *C. × paradisi* | PRJNA321100 | raw reads | Illumina | Wang et al., 2017 |
| | | Grapefruit | *C. × paradisi* | PRJNA321100 | raw reads | Illumina | Wang et al., 2017 |
| | | Grapefruit | *C. × paradisi 'Cocktail'* | PRJNA321100 | raw reads | Illumina | Wang et al., 2017 |
| | | Grapefruit cv. Marsh | *C. x paradisi Macfadyen* | PRJNA414519 | raw reads | Illumina | Wu et al., 2018 |
| | | star_ruby_grapefruit | *C. x paradisi* | PRJNA698060 | raw reads | BGISEQ500 | In this study |
| | | reed_white_grapefruit | *C. x paradisi* | PRJNA698060 | raw reads | BGISEQ500 | In this study |
| | | marsh_whitney_grapefruit_ | *C. x paradisi* | PRJNA698060 | raw reads | BGISEQ500 | In this study |
| | | marsh_brown_grapefruit | *C. x paradisi* | PRJNA698060 | raw reads | BGISEQ500 | In this study |
| | | oroblanco_grapefruit | *C. x paradisi* | PRJNA698060 | raw reads | BGISEQ500 | In this study |
| | | melogold_grapefruit | *C. x paradisi* | PRJNA698060 | raw reads | BGISEQ500 | In this study |
| | | melogold_grapefruit | *C. x paradisi* | PRJNA698060 | raw reads | BGISEQ500 | In this study |
| | | chironja_grapefruit | *C. x paradisi* | PRJNA698060 | raw reads | BGISEQ500 | In this study |

| | | | | | | | |
|---|---|---|---|---|---|---|---|
| | | star_ruby_grapefruit | *C. x paradisi* | PRJNA698060 | raw reads | BGISEQ500 | In this study |
| | | cocktail_grapefruit | *C. x paradisi* | PRJNA698060 | raw reads | BGISEQ500 | In this study |
| | | redblush_grapefruit | *C. x paradisi* | PRJNA698060 | raw reads | BGISEQ500 | In this study |
| | | oroblanco_grapefruit | *C. x paradisi* | PRJNA698060 | raw reads | BGISEQ500 | In this study |
| | | oroblanco_grapefruit | *C. x paradisi* | PRJNA698060 | raw reads | BGISEQ500 | In this study |
| | | rio_red_grapefruit | *C. x paradisi* | PRJNA698060 | raw reads | BGISEQ500 | In this study |
| | | rio_red_grapefruit | *C. x paradisi* | PRJNA698060 | raw reads | BGISEQ500 | In this study |
| Mandarin | | Clementine mandarin | *C. × clementina cv. Clemenules* | PRJNA225835 | High quality | Sanger | Wu et al., 2014 |
| | | Ponkan mandarin | *C. reticulata* | PRJNA225963 | raw reads | Illumina | Wu et al., 2014 |
| | | W. Murcott mandarin | *C. reticulata* | PRJNA225965 | raw reads | Illumina | Wu et al., 2014 |
| | | Willowleaf mandarin | *C. × deliciosa* | PRJNA225964 | raw reads | Illumina | Wu et al., 2014 |
| | | unnamed mandarin | *C. reticulata* | PRJNA320985 | raw reads | Illumina | Wang et al., 2017 |
| | | unnamed mandarin | *C. reticulata* | PRJNA320985 | raw reads | Illumina | Wang et al., 2017 |
| | | unnamed mandarin | *C. reticulata* | PRJNA320985 | raw reads | Illumina | Wang et al., 2017 |
| | | unnamed mandarin | *C. reticulata* | PRJNA320985 | raw reads | Illumina | Wang et al., 2017 |
| | | unnamed mandarin | *C. reticulata* | PRJNA320985 | raw reads | Illumina | Wang et al., 2017 |
| | | Mangshan mandarin | *C. reticulata* | PRJNA320985 | raw reads | Illumina | Wang et al., 2017 |
| | | Huanglingmiao mandarin | *C. reticulata* | PRJNA320985 | raw reads | Illumina | Wang et al., 2017 |
| | | Red tangerine | *C. reticulata* | PRJNA320985 | raw reads | Illumina | Wang et al., 2017 |
| | | Bingtangju | *C. reticulata* | PRJNA320985 | raw reads | Illumina | Wang et al., 2017 |
| | | Layueju | *C. reticulata* | PRJNA320985 | raw reads | Illumina | Wang et al., 2017 |
| | | Mashuiju | *C. reticulata* | PRJNA320985 | raw reads | Illumina | Wang et al., 2017 |

| | | | | | | | |
|---|---|---|---|---|---|---|---|
| | | unnamed mandarin | *C. reticulata* | PRJNA320985 | raw reads | Illumina | Wang et al., 2017 |
| | | unnamed mandarin | *C. reticulata* | PRJNA320985 | raw reads | Illumina | Wang et al., 2017 |
| | | Bendizao | *C. reticulata* | PRJNA320985 | raw reads | Illumina | Wang et al., 2018 |
| | | Changshananju | *C. reticulata* | PRJNA320985 | raw reads | Illumina | Wang et al., 2018 |
| | | Chongyi wild mandarin | *C. reticulata* | PRJNA320985 | raw reads | Illumina | Wang et al., 2018 |
| | | Chachigan | *C. reticulata cv. Chachiensis* | PRJNA320985 | raw reads | Illumina | Wang et al., 2018 |
| | | Ooita wase | *mandarin hybrid* | PRJNA320985 | raw reads | Illumina | Wang et al., 2018 |
| | | Daoxian wild mandarin No.1 | *C. reticulata* | PRJNA320985 | raw reads | Illumina | Wang et al., 2018 |
| | | Daoxian wild mandarin No.2 | *C. reticulata* | PRJNA320985 | raw reads | Illumina | Wang et al., 2018 |
| | | Daoxian wild mandarin No.3 | *C. reticulata* | PRJNA320985 | raw reads | Illumina | Wang et al., 2018 |
| | | Daoxian wild mandarin No.4 | *C. reticulata* | PRJNA320985 | raw reads | Illumina | Wang et al., 2018 |
| | | Huapiju | *C. reticulata* | PRJNA320985 | raw reads | Illumina | Wang et al., 2018 |
| | | Hezhou wild mandarin | *C. reticulata* | PRJNA320985 | raw reads | Illumina | Wang et al., 2018 |
| | | Jiangan | *C. reticulata* | PRJNA320985 | raw reads | Illumina | Wang et al., 2018 |
| | | Jiangyong wild mandarin | *C. reticulata* | PRJNA320985 | raw reads | Illumina | Wang et al., 2018 |
| | | Mingliutianju | *C. reticulata* | PRJNA320985 | raw reads | Illumina | Wang et al., 2018 |
| | | Mangshan wild mandarin | *C. reticulata* | PRJNA320985 | raw reads | Illumina | Wang et al., 2018 |
| | | Mangshan wild mandarin | *C. reticulata* | PRJNA320985 | raw reads | Illumina | Wang et al., 2018 |
| | | Mashuiju | *C. reticulata* | PRJNA320985 | raw reads | Illumina | Wang et al., 2018 |
| | | Nianju | *C. reticulata* | PRJNA320985 | raw reads | Illumina | Wang et al., 2018 |
| | | Orah | *mandarin hybrid* | PRJNA320985 | raw reads | Illumina | Wang et al., 2018 |
| | | Qingtianju | *C. reticulata* | PRJNA320985 | raw reads | Illumina | Wang et al., 2018 |

| | | | | | | | |
|---|---|---|---|---|---|---|---|
| | | Hongju | *C. reticulata* | PRJNA320985 | raw reads | Illumina | Wang et al., 2018 |
| | | Satsuma mandarin | *C. reticulata* | PRJNA320985 | raw reads | Illumina | Wang et al., 2018 |
| | | Sour tangerine | *C. reticulata* | PRJNA320985 | raw reads | Illumina | Wang et al., 2018 |
| | | Suanpangan | *C. reticulata* | PRJNA320985 | raw reads | Illumina | Wang et al., 2018 |
| | | Shatangju | *C. reticulata* | PRJNA320985 | raw reads | Illumina | Wang et al., 2018 |
| | | Seedless Ponkan | *C. reticulata* | PRJNA320985 | raw reads | Illumina | Wang et al., 2018 |
| | | Wilking | *mandarin hybrid* | PRJNA320985 | raw reads | Illumina | Wang et al., 2018 |
| | | Owari unshiu | *mandarin hybrid* | PRJNA320985 | raw reads | Illumina | Wang et al., 2018 |
| | | Yuanjiangnanju | *C. reticulata* | PRJNA320985 | raw reads | Illumina | Wang et al., 2018 |
| | | Yangshanju | *C. reticulata* | PRJNA320985 | raw reads | Illumina | Wang et al., 2018 |
| | | Zhuhongju | *C. reticulata* | PRJNA320985 | raw reads | Illumina | Wang et al., 2018 |
| | | Tachibana mandarin | *C. tachibana (Mak)* | PRJNA320985 | raw reads | Illumina | Wu et al., 2018 |
| | | Sunki mandarin | *C. x sunki (hayata)* | PRJNA414519 | raw reads | Illumina | Wu et al., 2018 |
| | | Dancy mandarin | *C. x tangerina (Tanaka)* | PRJNA414519 | raw reads | Illumina | Wu et al., 2018 |
| | | Satsuma mandarin | *C. x unshiu (Marc)* | PRJNA414519 | raw reads | Illumina | Wu et al., 2018 |
| | | Cleopatra mandarin | *C. x reshni (Hort. ex Tanaka)* | PRJNA414519 | raw reads | Illumina | Wu et al., 2018 |
| | | Changsha mandarin | *C. x reticulata (Blanco)* | PRJNA414519 | raw reads | Illumina | Wu et al., 2018 |
| | | Kishu mandarin | *C. x kinokuni (Hort. ex Tanaka)* | PRJNA414519 | raw reads | Illumina | Wu et al., 2018 |
| | | Sun Chu Sha Ka | *C. reticulata (Blanco)* | PRJNA414519 | raw reads | Illumina | Wu et al., 2018 |
| | | *C. mangshanensis* | *C. mangshanensis* | PRJNA392792 | raw reads | Illumina | Wang et al., 2018 |
| | | Yuanju | *N.D.* | PRJNA392792 | raw reads | Illumina | Wang et al., 2018 |
| | | Chazhigan | *Citrus haniana Hort. ex Tseng* | http://citgvd.cric.cn/home/index | raw reads | Illumina | Li et al., 2020 |

| | | | | | | | |
|---|---|---|---|---|---|---|---|
| | | Shangtangju | *Citrus sp.* | http://citgvd.cric.cn/home/index | raw reads | Illumina | Li et al., 2020 |
| | | Shiyueju | *Citrus flamea hort.ex Tseng* | http://citgvd.cric.cn/home/index | raw reads | Illumina | Li et al., 2020 |
| | | Dakengyeju | *Citrus sp.* | http://citgvd.cric.cn/home/index | raw reads | Illumina | Li et al., 2020 |
| | | Daoxianensis | *Citrus sp.* | http://citgvd.cric.cn/home/index | raw reads | Illumina | Li et al., 2020 |
| | | Cupigoushigan | *Citrus nobilis Lour* | http://citgvd.cric.cn/home/index | raw reads | Illumina | Li et al., 2020 |
| | | Xipigoushigan | *Citrus nobilis Lour* | http://citgvd.cric.cn/home/index | raw reads | Illumina | Li et al., 2020 |
| | | Yinduyeju | *Citrus haniana Hort. ex Tseng* | http://citgvd.cric.cn/home/index | raw reads | Illumina | Li et al., 2020 |
| | | Makino | *Citrus tachibana Tanaka* | http://citgvd.cric.cn/home/index | raw reads | Illumina | Li et al., 2020 |
| | | Jiaogan | *Citrus nobilis Lour* | http://citgvd.cric.cn/home/index | raw reads | Illumina | Li et al., 2020 |
| | | Daxianggan | *Citrus haniana Hort. ex Tseng* | http://citgvd.cric.cn/home/index | raw reads | Illumina | Li et al., 2020 |
| | | Xingyidahongpao | *Citrus tangerine Tanaka* | http://citgvd.cric.cn/home/index | raw reads | Illumina | Li et al., 2020 |
| | | Yuanjiangjianggan | *Citrus grandis (L) Osbeck* | http://citgvd.cric.cn/home/index | raw reads | Illumina | Li et al., 2020 |
| | | Nanfengmiju | *Citrus haniana Hort. ex Tseng* | http://citgvd.cric.cn/home/index | raw reads | Illumina | Li et al., 2020 |
| | | Ougan | *Citrus speciosa Hort. ex Tsen* | http://citgvd.cric.cn/home/index | raw reads | Illumina | Li et al., 2020 |
| | | Zhoupigan | *Citrus speciosa Hort. ex Tsen* | http://citgvd.cric.cn/home/index | raw reads | Illumina | Li et al., 2020 |
| | | Huangpisuanju | *Citrus haniana Hort. ex Tseng* | http://citgvd.cric.cn/home/index | raw reads | Illumina | Li et al., 2020 |
| | | Zhuhongju | *Citrus erythrosa* | http://citgvd.cric.cn/home/index | raw reads | Illumina | Li et al., 2020 |
| | | Xinshengxi No. 3 | *Citrus sp.* | http://citgvd.cric.cn/home/index | raw reads | Illumina | Li et al., 2020 |
| | | Yanxiwanlu | *Citrus retlculata Blanco* | http://citgvd.cric.cn/home/index | raw reads | Illumina | Li et al., 2020 |
| | | Shengtian | *Citrus sp.* | http://citgvd.cric.cn/home/index | raw reads | Illumina | Li et al., 2020 |
| | | Gongchuan | *Citrus sp.* | http://citgvd.cric.cn/home/index | raw reads | Illumina | Li et al., 2020 |
| | | Mangshanyeju | *Citrus nobilis Lour* | http://citgvd.cric.cn/home/index | raw reads | Illumina | Li et al., 2020 |

| | | | | | | | |
|---|---|---|---|---|---|---|---|
| | | fairchild_mandarin | *C. reticulata* | PRJNA698060 | raw reads | BGISEQ500 | In this study |
| | | daisy_mandarin | *C. reticulata* | PRJNA698060 | raw reads | BGISEQ500 | In this study |
| | | tango_mandarin | *C. reticulata* | PRJNA698060 | raw reads | BGISEQ500 | In this study |
| | | ellendale_mandarin | *C. reticulata* | PRJNA698060 | raw reads | BGISEQ500 | In this study |
| | | cleopatra | *C. reticulata* | PRJNA698060 | raw reads | BGISEQ500 | In this study |
| | | citrus_tangerina | *C. reticulata* | PRJNA698060 | raw reads | BGISEQ500 | In this study |
| | | Tachibana_orange | *C. tachibana* | PRJNA698060 | raw reads | BGISEQ500 | In this study |
| Pummelo | | Shatian pummelo | *C. grandis* | PRJNA318855 | raw reads | Illumina | Xu et al., 2013 |
| | | Wusuan pummelo | *C. grandis* | PRJNA318855 | raw reads | Illumina | Xu et al., 2013 |
| | | Guanxi pummelo | *C. grandis* | PRJNA318855 | raw reads | Illumina | Xu et al., 2013 |
| | | Chandler pummelo | *C. grandis* | SRX372688 | raw reads | Illumina | Wu et al., 2018 |
| | | Low-acid pummelo | *C. grandis* | SRX372702 | raw reads | Illumina | Wu et al., 2018 |
| | | Anjianghonxinyou | *C. grandis* | PRJNA318855 | raw reads | Illumina | Wang et al., 2018 |
| | | Hongxinyou from Huaihua, Hunan | *C. grandis* | PRJNA318855 | raw reads | Illumina | Wang et al., 2018 |
| | | Shawanyou from Huaihua, Hunan | *C. grandis* | PRJNA318855 | raw reads | Illumina | Wang et al., 2018 |
| | | Guanximiyou | *C. grandis* | PRJNA318855 | raw reads | Illumina | Wang et al., 2018 |
| | | Majiayou | *C. grandis* | PRJNA318855 | raw reads | Illumina | Wang et al., 2017 |
| | | pummelo from Guilin No.1 | *C. grandis* | PRJNA318855 | raw reads | Illumina | Wang et al., 2017 |
| | | pummelo from Chongqing No.016 | *C. grandis* | PRJNA318855 | raw reads | Illumina | Wang et al., 2017 |
| | | pummelo from Ji'an, jiangxi No.02 | *C. grandis* | PRJNA318855 | raw reads | Illumina | Wang et al., 2017 |
| | | Huazhoujuhong | *C. grandis* | PRJNA318855 | raw reads | Illumina | Wang et al., 2017 |
| | | Huanonghongyou | *C. grandis* | PRJNA318855 | raw reads | Illumina | Wang et al., 2017 |

| | | | | | | | |
|---|---|---|---|---|---|---|---|
| | | Wanbaiyou | *C. grandis* | PRJNA318855 | raw reads | Illumina | Wang et al., 2017 |
| | | pummelo from Ruili, Yunnan No.6 | *C. grandis* | PRJNA318855 | raw reads | Illumina | Wang et al., 2017 |
| | | Kaopan | *C. grandis* | PRJNA318855 | raw reads | Illumina | Wang et al., 2017 |
| | | pummelo from Chongqing No.04 | *C. grandis* | PRJNA318855 | raw reads | Illumina | Wang et al., 2017 |
| | | Pummelo | *C. grandis* | PRJNA318855 | raw reads | Illumina | Wang et al., 2017 |
| | | Pummelo | *C. grandis* | PRJNA318855 | raw reads | Illumina | Wang et al., 2017 |
| | | Pummelo | *C. grandis* | PRJNA318855 | raw reads | Illumina | Wang et al., 2017 |
| | | Pummelo | *C. grandis* | PRJNA318855 | raw reads | Illumina | Wang et al., 2017 |
| | | Pummelo | *C. grandis* | PRJNA318855 | raw reads | Illumina | Wang et al., 2017 |
| | | Zipiyou | *C. grandis* | PRJNA318855 | raw reads | Illumina | Wang et al., 2017 |
| | | Pushi sour pummelo | *C. grandis* | PRJNA318855 | raw reads | Illumina | Wang et al., 2017 |
| | | Sour pummelo in Baishazhen | *C. grandis* | PRJNA318855 | raw reads | Illumina | Wang et al., 2017 |
| | | local pummelo in Hunan | *C. grandis* | PRJNA318855 | raw reads | Illumina | Wang et al., 2017 |
| | | pummelo from Guilin No.01 | *C. grandis* | PRJNA318855 | raw reads | Illumina | Wang et al., 2017 |
| | | Pummelo | *C. grandis* | PRJNA318855 | High quality assemblied genomes | PacBio RS II/Illumina | Wang et al., 2017 |
| | | You 8020 | *Citrus grandis (L) Osbeck* | http://citgvd.cric.cn/home/index | raw reads | Illumina | Li et al., 2020 |
| | | You 8088 | *Citrus grandis (L) Osbeck* | http://citgvd.cric.cn/home/index | raw reads | Illumina | Li et al., 2020 |
| | | Anjianghongxinyou | *Citrus grandis (L) Osbeck* | http://citgvd.cric.cn/home/index | raw reads | Illumina | Li et al., 2020 |
| | | Anjiangshiliuyou | *Citrus grandis (L) Osbeck* | http://citgvd.cric.cn/home/index | raw reads | Illumina | Li et al., 2020 |
| | | Anjiangwuheyou | *Citrus grandis (L) Osbeck* | http://citgvd.cric.cn/home/index | raw reads | Illumina | Li et al., 2020 |
| | | Anjiangxiangyou | *Citrus grandis (L)Osbeck* | http://citgvd.cric.cn/home/index | raw reads | Illumina | Li et al., 2020 |

| | | | | | | | |
|---|---|---|---|---|---|---|---|
| | | Anlong No. 1 | *Citrus grandis (L) Osbeck* | http://citgvd.cric.cn/home/index | raw reads | Illumina | Li et al., 2020 |
| | | Bairoupaoguo | *Citrus grandis (L) Osbeck* | http://citgvd.cric.cn/home/index | raw reads | Illumina | Li et al., 2020 |
| | | Baiyushuang | *Citrus grandis (L) Osbeck* | http://citgvd.cric.cn/home/index | raw reads | Illumina | Li et al., 2020 |
| | | Beibeiyou | *Citrus grandis (L) Osbeck* | http://citgvd.cric.cn/home/index | raw reads | Illumina | Li et al., 2020 |
| | | Bingtangyou | *Citrus grandis (L) Osbeck* | http://citgvd.cric.cn/home/index | raw reads | Illumina | Li et al., 2020 |
| | | Chumenwendan | *Citrus grandis (L) Osbeck* | http://citgvd.cric.cn/home/index | raw reads | Illumina | Li et al., 2020 |
| | | Chuhongyou | *Citrus grandis (L) Osbeck* | http://citgvd.cric.cn/home/index | raw reads | Illumina | Li et al., 2020 |
| | | Cuixiangtianyou | *Citrus grandis (L) Osbeck* | http://citgvd.cric.cn/home/index | raw reads | Illumina | Li et al., 2020 |
| | | Dapaoguo | *Citrus grandis (L) Osbeck* | http://citgvd.cric.cn/home/index | raw reads | Illumina | Li et al., 2020 |
| | | Dayongjuhuaxinyou | *Citrus grandis (L) Osbeck* | http://citgvd.cric.cn/home/index | raw reads | Illumina | Li et al., 2020 |
| | | Dianjiangbaiyou | *Citrus grandis (L) Osbeck* | http://citgvd.cric.cn/home/index | raw reads | Illumina | Li et al., 2020 |
| | | Diangjianghongxinyou | *Citrus grandis (L) Osbeck* | http://citgvd.cric.cn/home/index | raw reads | Illumina | Li et al., 2020 |
| Papeda | Resistant/ Tolerant | Papeda | *Citrus ichangensis* | PRJNA321657 | Draft assemblied genome | Illumina | Wang et al., 2017 |
| | | wild Ichang papeda XJC | *C. ichangensis* | PRJNA321657 | raw reads | Illumina | Wang et al., 2017 |
| | | wild Ichang papeda XJC | *C. ichangensis* | PRJNA321657 | raw reads | Illumina | Wang et al., 2017 |
| | | wild Ichang papeda XJC | *C. ichangensis* | PRJNA321657 | raw reads | Illumina | Wang et al., 2017 |
| | | wild Ichang papeda XJC | *C. ichangensis* | PRJNA321657 | raw reads | Illumina | Wang et al., 2017 |
| | | wild Ichang papeda XJC | *C. ichangensis* | PRJNA321657 | raw reads | Illumina | Wang et al., 2017 |
| | | wild Ichang papeda PY11 | *C. ichangensis* | PRJNA321657 | raw reads | Illumina | Wang et al., 2017 |
| | | wild Ichang papeda YCC | *C. ichangensis* | PRJNA321657 | raw reads | Illumina | Wang et al., 2017 |
| | | wild Ichang papeda DYC | *C. ichangensis* | PRJNA321657 | raw reads | Illumina | Wang et al., 2017 |
| | | wild Ichang papeda JF | *C. ichangensis* | PRJNA321657 | raw reads | Illumina | Wang et al., 2017 |

| | | | | | | | |
|---|---|---|---|---|---|---|---|
| | | wild Ichang papeda KM | *C. ichangensis* | PRJNA321657 | raw reads | Illumina | Wang et al., 2017 |
| | | wild Ichang papeda YCLS | *C. ichangensis* | PRJNA321657 | raw reads | Illumina | Wang et al., 2017 |
| | | wild Ichang papeda TK | *C. ichangensis* | PRJNA321657 | raw reads | Illumina | Wang et al., 2017 |
| | | wild Ichang papeda YCYJ | *C. ichangensis* | PRJNA321657 | raw reads | Illumina | Wang et al., 2017 |
| | | wild Ichang papeda YL | *C. ichangensis* | PRJNA321657 | raw reads | Illumina | Wang et al., 2017 |
| | | wild Ichang papeda ZY | *C. ichangensis* | PRJNA321657 | raw reads | Illumina | Wang et al., 2017 |
| | | wild Ichang papeda XC | *C. ichangensis* | PRJNA321657 | raw reads | Illumina | Wang et al., 2017 |
| | | Ichang papeda | *C. ichangensis (Swingle)* | PRJNA414519 | raw reads | Illumina | Wu et al., 2018 |
| | | Yuanjiangyichangcheng | *Citrus ichangensis* | http://citgvd.cric.cn/home/index | raw reads | Illumina | Li et al., 2020 |
| | | Jindaoxiayichangcheng | *Citrus ichangensis* | http://citgvd.cric.cn/home/index | raw reads | Illumina | Li et al., 2020 |
| | | Ichangensis 2586 | *Citrus ichangensis Swingle* | http://citgvd.cric.cn/home/index | raw reads | Illumina | Li et al., 2020 |
| | | Hongheensis | *Citrus sp.* | http://citgvd.cric.cn/home/index | raw reads | Illumina | Li et al., 2020 |
| | | Montrous | *Citrus macroptera* | http://citgvd.cric.cn/home/index | raw reads | Illumina | Li et al., 2020 |
| | | Turpinia ternata Nakai | *Citrus sp.* | http://citgvd.cric.cn/home/index | raw reads | Illumina | Li et al., 2020 |
| | | Zhique | *Citrus wilsonii Tanaka* | http://citgvd.cric.cn/home/index | raw reads | Illumina | Li et al., 2020 |
| | | Ziyangxiangcheng | *Citrus sp.* | http://citgvd.cric.cn/home/index | raw reads | Illumina | Li et al., 2020 |
| | | Xiecheng | *Citrus junons Sieb. ex. Tanaka* | http://citgvd.cric.cn/home/index | raw reads | Illumina | Li et al., 2020 |
| | | Zhencheng | *Citrus sp.* | http://citgvd.cric.cn/home/index | raw reads | Illumina | Li et al., 2020 |
| | | Dazhongcheng | *Citrus sp.* | http://citgvd.cric.cn/home/index | raw reads | Illumina | Li et al., 2020 |
| | | macrophylla | *C. ichangensis* | PRJNA698060 | raw reads | BGISEQ500 | In this study |
| | | macrophylla | *C. ichangensis* | PRJNA698060 | raw reads | BGISEQ500 | In this study |
| | | macrophylla | *C. ichangensis* | PRJNA698060 | raw reads | BGISEQ500 | In this study |

| | | | | | | | |
|---|---|---|---|---|---|---|---|
| Kumquat | Resistant/ Tolerant | Oval kumquat | *Fortunella margarita (Lour.)* | PRJNA414519 | raw reads | Illumina | Wu et al., 2018 |
| | | Hongkong kumquat | *Fortunella hindsii* | PRJNA487160 | High quality assemblied genomes | PacBio/Illumina | Zhu et al., 2019 |
| | | Hongkong kumquat | *Fortunella hindsii* | PRJNA487160 | raw reads | Illumina | Zhu et al., 2019 |
| | | Hongkong kumquat | *Fortunella hindsii* | PRJNA487160 | raw reads | Illumina | Zhu et al., 2019 |
| | | Daguojindou | *Fortunella venosa* | http://citgvd.cric.cn/home/index | raw reads | Illumina | Li et al., 2020 |
| | | Hongkong kumquat | *Fortunella hindsii* | http://citgvd.cric.cn/home/index | raw reads | Illumina | Li et al., 2020 |
| | | Meiwa kumquat | *Fortunella crassifiolia Swingle* | http://citgvd.cric.cn/home/index | raw reads | Illumina | Li et al., 2020 |
| | | Meiwa kumquat | *Fortunella crassifiolia Swingle* | http://citgvd.cric.cn/home/index | raw reads | Illumina | Li et al., 2020 |
| | | Meiwa kumquat | *Fortunella crassifiolia Swingle* | http://citgvd.cric.cn/home/index | raw reads | Illumina | Li et al., 2020 |
| | | Meiwa kumquat | *Fortunella crassifiolia Swingle* | PRJNA698060 | raw reads | BGISEQ500 | In this study |
| Kumquat hybirds | | limequat | *C. × floridana* | PRJNA698060 | raw reads | BGISEQ500 | In this study |
| | | calamondin_lime | *C. unshiu x F. japonica* | PRJNA698060 | raw reads | BGISEQ500 | In this study |
| | | Calamondin | *C. x madurensis (Lour.)* | PRJNA414519 | raw reads | Illumina | Wu et al., 2018 |
| Atalantia | susceptible | Chinese Box Orange | *Atlantia buxifolia* | PRJNA327148 | Draft assemblied genome | Illumina | Wang et al., 2017 |
| | | Chinese box orange | *A. buxifolia* | PRJNA327148 | raw reads | Illumina | Wang et al., 2017 |
| | | Chinese box orange | *A. buxifolia* | PRJNA327148 | raw reads | Illumina | Wang et al., 2017 |
| | | Chinese box orange | *A. buxifolia* | PRJNA327148 | raw reads | Illumina | Wang et al., 2017 |
| | | Chinese box orange | *A. buxifolia* | PRJNA327148 | raw reads | Illumina | Wang et al., 2017 |
| | | Chinese box orange | *A. buxifolia* | PRJNA327148 | raw reads | Illumina | Wang et al., 2017 |
| | | Chinese box orange | *A. buxifolia* | PRJNA327148 | raw reads | Illumina | Wang et al., 2017 |
| | | Chinese box orange | *A. buxifolia* | PRJNA327148 | raw reads | Illumina | Wang et al., 2017 |

| | | | | | | | |
|---|---|---|---|---|---|---|---|
| | | Chinese box orange | *A. buxifolia* | PRJNA327148 | raw reads | Illumina | Wang et al., 2017 |
| | | Chinese box orange | *A. buxifolia* | PRJNA327148 | raw reads | Illumina | Wang et al., 2017 |
| | | Chinese box orange | *A. buxifolia* | PRJNA327148 | raw reads | Illumina | Wang et al., 2017 |
| | | Chinese box orange | *A. buxifolia* | PRJNA327148 | raw reads | Illumina | Wang et al., 2017 |
| | | Chinese box orange | *A. buxifolia* | PRJNA327148 | raw reads | Illumina | Wang et al., 2017 |
| | | Chinese box orange | *A. buxifolia* | PRJNA327148 | raw reads | Illumina | Wang et al., 2017 |
| | | Chinese box orange | *A. buxifolia* | PRJNA327148 | raw reads | Illumina | Wang et al., 2017 |
| | | Chinese box orange | *A. buxifolia* | PRJNA414519 | raw reads | Illumina | Wu et al., 2018 |
| Trifoliate orange | Resistant/ Tolerant | Trifoliate orange | *Poncirus trifoliata (L.) Raf* | PRJNA414519 | raw reads | Illumina | Wu et al., 2018 |
| | | Xiaoyezhi | *Poncirus trifoliata (L.) Raf)* | http://citgvd.cric.cn/home/index | raw reads | Illumina | Li et al., 2020 |
| | | Xiaohuazhi No. 5 | *Poncirus trifoliata (L.) Raf)* | http://citgvd.cric.cn/home/index | raw reads | Illumina | Li et al., 2020 |
| | | Dayezhi | *Poncirus trifoliata (L.) Raf)* | http://citgvd.cric.cn/home/index | raw reads | Illumina | Li et al., 2020 |
| | | Yanlingzhicheng | *Poncirus trifoliata (L.) Raf* | http://citgvd.cric.cn/home/index | raw reads | Illumina | Li et al., 2020 |
| | | Trifoliate orange | *Poncirus trifoliata (L.) Raf* | PRJNA648176 | High quality assemblied genomes | Illumina/PacBio/Hi-C | Peng et al., 2020 |
| | | Flying Dragon | *Poncirus trifoliata (L.) Raf* | PRJNA648176 | raw reads | Illumina | Peng et al., 2020 |
| | | Little Leaf | *Poncirus trifoliata (L.) Raf* | PRJNA648176 | raw reads | Illumina | Peng et al., 2020 |
| | | Rubidoux | *Poncirus trifoliata (L.) Raf* | PRJNA648176 | raw reads | Illumina | Peng et al., 2020 |
| | | Poncirus polyandra | *Poncirus polyandra* | PRJNA648176 | raw reads | Illumina | Peng et al., 2020 |
| Poncirus hybirds | | Carrizo citrange | *Citrus x sinensis x Poncirus trifoliata* | PRJNA648176 | raw reads | Illumina | Peng et al., 2020 |
| | | C-35 | *Citrus x sinensis x Poncirus trifoliata* | PRJNA698060 | raw reads | Illumina | In this study |
| | | carrizo | *Citrus x sinensis x Poncirus trifoliata* | PRJNA698060 | raw reads | BGISEQ500 | In this study |

| | | | | | | | |
|---|---|---|---|---|---|---|---|
| | | carrizo | *Citrus x sinensis x Poncirus trifoliata* | PRJNA698060 | raw reads | BGISEQ500 | In this study |
| | | carrizo | *Citrus x sinensis x Poncirus trifoliata* | PRJNA698060 | raw reads | BGISEQ500 | In this study |
| | | carrizo | *Citrus x sinensis x Poncirus trifoliata* | PRJNA698060 | raw reads | BGISEQ500 | In this study |
| | | carrizo | *Citrus x sinensis x Poncirus trifoliata* | PRJNA698060 | raw reads | BGISEQ500 | In this study |
| | | carrizo | *Citrus x sinensis x Poncirus trifoliata* | PRJNA698060 | raw reads | BGISEQ500 | In this study |
| | | carrizo | *Citrus x sinensis x Poncirus trifoliata* | PRJNA698060 | raw reads | BGISEQ500 | In this study |
| | | carrizo | *Citrus x sinensis x Poncirus trifoliata* | PRJNA698060 | raw reads | BGISEQ500 | In this study |
| | | troyer | *Citrus x sinensis x Poncirus trifoliata* | PRJNA698060 | raw reads | BGISEQ500 | In this study |
| | | carrizo | *Citrus x sinensis x Poncirus trifoliata* | PRJNA698060 | raw reads | BGISEQ500 | In this study |
| | | carrizo | *Citrus x sinensis x Poncirus trifoliata* | PRJNA698060 | raw reads | BGISEQ500 | In this study |
| | | carrizo | *Citrus x sinensis x Poncirus trifoliata* | PRJNA698060 | raw reads | BGISEQ500 | In this study |
| | | carrizo | *Citrus x sinensis x Poncirus trifoliata* | PRJNA698060 | raw reads | BGISEQ500 | In this study |
| | | C-35 | *Citrus x sinensis x Poncirus trifoliata* | PRJNA698060 | raw reads | BGISEQ500 | In this study |
| | | troyer | *Citrus x sinensis x Poncirus trifoliata* | PRJNA698060 | raw reads | BGISEQ500 | In this study |
| | | carrizo | *Citrus x sinensis x Poncirus trifoliata* | PRJNA698060 | raw reads | BGISEQ500 | In this study |
| | | carrizo | *Citrus x sinensis x Poncirus trifoliata* | PRJNA698060 | raw reads | BGISEQ500 | In this study |
| | | carrizo | *Citrus x sinensis x Poncirus trifoliata* | PRJNA698060 | raw reads | BGISEQ500 | In this study |
| | | carrizo | *Citrus x sinensis x Poncirus trifoliata* | PRJNA698060 | raw reads | BGISEQ500 | In this study |
| | | carrizo | *Citrus x sinensis x Poncirus trifoliata* | PRJNA698060 | raw reads | BGISEQ500 | In this study |
| | | C-35 | *Citrus x sinensis x Poncirus trifoliata* | PRJNA698060 | raw reads | BGISEQ500 | In this study |
| | | carrizo | *Citrus x sinensis x Poncirus trifoliata* | PRJNA698060 | raw reads | BGISEQ500 | In this study |
| | | carrizo | *Citrus x sinensis x Poncirus trifoliata* | PRJNA698060 | raw reads | BGISEQ500 | In this study |

| | | | | | | | |
|---|---|---|---|---|---|---|---|
| | | carrizo | *Citrus x sinensis x Poncirus trifoliata* | PRJNA698060 | raw reads | BGISEQ500 | In this study |
| | | carrizo | *Citrus x sinensis x Poncirus trifoliata* | PRJNA698060 | raw reads | BGISEQ500 | In this study |
| | | carrizo | *Citrus x sinensis x Poncirus trifoliata* | PRJNA698060 | raw reads | BGISEQ500 | In this study |
| | | carrizo | *Citrus x sinensis x Poncirus trifoliata* | PRJNA698060 | raw reads | BGISEQ500 | In this study |
| | | carrizo | *Citrus x sinensis x Poncirus trifoliata* | PRJNA698060 | raw reads | BGISEQ500 | In this study |
| | | carrizo | *Citrus x sinensis x Poncirus trifoliata* | PRJNA698060 | raw reads | BGISEQ500 | In this study |
| | | carrizo | *Citrus x sinensis x Poncirus trifoliata* | PRJNA698060 | raw reads | BGISEQ500 | In this study |
| | | carrizo | *Citrus x sinensis x Poncirus trifoliata* | PRJNA698060 | raw reads | BGISEQ500 | In this study |
| | | C-35 | *Citrus x sinensis x Poncirus trifoliata* | PRJNA698060 | raw reads | BGISEQ500 | In this study |
| | | carrizo | *Citrus x sinensis x Poncirus trifoliata* | PRJNA698060 | raw reads | BGISEQ500 | In this study |
| | | C-35 | *Citrus x sinensis x Poncirus trifoliata* | PRJNA698060 | raw reads | BGISEQ500 | In this study |
| | | carrizo | *Citrus x sinensis x Poncirus trifoliata* | PRJNA698060 | raw reads | BGISEQ500 | In this study |
| | | carrizo | *Citrus x sinensis x Poncirus trifoliata* | PRJNA698060 | raw reads | BGISEQ500 | In this study |
| | | carrizo | *Citrus x sinensis x Poncirus trifoliata* | PRJNA698060 | raw reads | BGISEQ500 | In this study |
| | | carrizo | *Citrus x sinensis x Poncirus trifoliata* | PRJNA698060 | raw reads | BGISEQ500 | In this study |
| | | carrizo | *Citrus x sinensis x Poncirus trifoliata* | PRJNA698060 | raw reads | BGISEQ500 | In this study |
| | | carrizo | *Citrus x sinensis x Poncirus trifoliata* | PRJNA698060 | raw reads | BGISEQ500 | In this study |
| | | carrizo | *Citrus x sinensis x Poncirus trifoliata* | PRJNA698060 | raw reads | BGISEQ500 | In this study |
| | | C-35 | *Citrus x sinensis x Poncirus trifoliata* | PRJNA698060 | raw reads | BGISEQ500 | In this study |
| | | carrizo | *Citrus x sinensis x Poncirus trifoliata* | PRJNA698060 | raw reads | BGISEQ500 | In this study |
| | | carrizo | *Citrus x sinensis x Poncirus trifoliata* | PRJNA698060 | raw reads | BGISEQ500 | In this study |
| | | C-35 | *Citrus x sinensis x Poncirus trifoliata* | PRJNA698060 | raw reads | BGISEQ500 | In this study |

| | | | | | | | |
|---|---|---|---|---|---|---|---|
| | | C-35 | *Citrus x sinensis x Poncirus trifoliata* | PRJNA698060 | raw reads | BGISEQ500 | In this study |
| | | Xcitroncirus X639 | *Citrus reticulata x Poncirus trifoliata* | PRJNA648176 | raw reads | Illumina | Peng et al., 2020 |
| | | US-897 | *Citrus reticulata x Poncirus trifoliata* | PRJNA648176 | raw reads | Illumina | Peng et al., 2020 |
| | | US-812 | *Citrus reticulata x Poncirus trifoliata* | PRJNA648176 | raw reads | Illumina | Peng et al., 2020 |
| | | US-942 | *Citrus reticulata x Poncirus trifoliata* | PRJNA648176 | raw reads | Illumina | Peng et al., 2020 |
| | | US802 | *Citrus grandis x Poncirus trifoliata* | PRJNA648176 | raw reads | Illumina | Peng et al., 2020 |
| | | citrumelo | *Citrus paradisi x Poncirus trifoliata* | PRJNA698060 | raw reads | BGISEQ500 | In this study |
| Clausena | Resistant/ Tolerant | Clausena | *Clausena lansium* | PRJNA392722 | raw reads | Illumina | Wang et al., 2018 |
| | | Clausena lansium | *Clausena lansium (Lour.) Skeels* | http://citgvd.cric.cn/home/index | raw reads | Illumina | Li et al., 2020 |
| Murraya | | Murraya exotica | *Murraya exotica* | http://citgvd.cric.cn/home/index | raw reads | Illumina | Li et al., 2020 |

**Table S21. The summary of newly sequenced genomic data for GWAS study**

| Sample | Common name | Group name (Scientific name) | Clean data (Gb) | Sequencing depth for citrus genome |
|---|---|---|---|---|
| L12 | citrumelo_4475 | Citrumelo (*X Citroncirus spp.*) | 43.47 | 111 |
| L105 | star_ruby_grapefruit | Grapefruit (*C. x paradisi*) | 40.48 | 104 |
| L106 | reed_white_grapefruit | Grapefruit (*C. x paradisi*) | 46.72 | 120 |
| L107 | marsh_whitney_grapefruit_ | Grapefruit (*C. x paradisi*) | 37.70 | 97 |
| L108 | marsh_brown_grapefruit | Grapefruit (*C. x paradisi*) | 41.97 | 108 |
| L109 | oroblanco_grapefruit | Grapefruit (*C. x paradisi*) | 46.06 | 118 |
| L110 | melogold_grapefruit | Grapefruit (*C. x paradisi*) | 44.46 | 114 |
| L111 | melogold_grapefruit | Grapefruit (*C. x paradisi*) | 39.20 | 101 |
| L112 | chironja_grapefruit | Grapefruit (*C. x paradisi*) | 44.55 | 114 |
| L113 | star_ruby_grapefruit | Grapefruit (*C. x paradisi*) | 43.34 | 111 |
| L114 | cocktail_grapefruit | Grapefruit (*C. x paradisi*) | 42.46 | 109 |

| | | | | |
|---|---|---|---|---|
| L115 | redblush_grapefruit | Grapefruit (*C. x paradisi*) | 33.64 | 86 |
| L123 | oroblanco_grapefruit | Grapefruit (*C. x paradisi*) | 5.19 | 13 |
| L124 | oroblanco_grapefruit | Grapefruit (*C. x paradisi*) | 35.15 | 90 |
| L125 | rio_red_grapefruit | Grapefruit (*C. x paradisi*) | 43.97 | 113 |
| L126 | rio_red_grapefruit | Grapefruit (*C. x paradisi*) | 39.62 | 102 |
| L6 | citrus_hystrix | Lime (*C. hystrix*) | 47.60 | 122 |
| L61 | improved_meyer_lemon | Lemon (*Citrus x meyeri 'Improved'*) | 22.52 | 58 |
| L66 | ponderosa_lemon | Ponderosa_lemon (*C. x pyriformis*) | 70.20 | 193 |
| L69 | mexican_lime | Lime (*Citrus x aurantifolia*) | 51.22 | 131 |
| L94 | meiwa_kumquat | Kumquat (*Fortunella crassifolia*) | 46.11 | 118 |
| L88 | limequat | Limequat (C. × floridana) | 31.51 | 81 |
| L78 | calamondin_lime | Orangequat (*C. unshiu x C. japonica*) | 41.14 | 105 |
| L83 | fairchild_mandarin | Mandarin (*C. reticulata*) | 51.13 | 131 |
| L86 | daisy_mandarin | Mandarin (*C. reticulata*) | 133.00 | 341 |
| L93 | tango_mandarin | Mandarin (*C. reticulata*) | 59.42 | 152 |
| L102 | C.tachibana_ | Tachibana_orange (*C. tachibana*) | 40.27 | 103 |
| L99 | ellendale_mandarin | Mandarin (*C. reticulata*) | 45.21 | 116 |
| R106 | carrizo | Citrange (*X Citroncirus sp.*) | 7.34 | 19 |
| R109 | carrizo | Citrange (*X Citroncirus sp.*) | 15.98 | 41 |
| R110 | citrumelo | Citrumelo (*X Citroncirus spp.*) | 12.62 | 32 |
| R114 | carrizo | Citrange (*X Citroncirus sp.*) | 12.90 | 33 |
| R124 | C-35 | Citrange (*X Citroncirus sp.*) | 15.58 | 40 |
| R126 | troyer | Citrange (*X Citroncirus sp.*) | 8.76 | 22 |
| R34 | volckameriana | Lemon (*C. × limon*) | 12.21 | 31 |
| R35 | carrizo | Citrange (*X Citroncirus sp.*) | 9.17 | 24 |
| R48 | carrizo | Citrange (*X Citroncirus sp.*) | 11.39 | 29 |
| R49 | carrizo | Citrange (*X Citroncirus sp.*) | 15.78 | 40 |
| R50 | C-35 | Citrange (*X Citroncirus sp.*) | 15.71 | 40 |
| R5 | carrizo | Citrange (*X Citroncirus sp.*) | 12.85 | 33 |

| | | | | |
|---|---|---|---|---|
| R86 | carrizo | Citrange (*X Citroncirus sp.*) | 17.19 | 44 |
| R90 | C-35 | Citrange (*X Citroncirus sp.*) | 12.05 | 31 |
| R104 | volckameriana | Lemon (*C.* × *limon*) | 13.93 | 36 |
| R105 | carrizo | Citrange (*X Citroncirus sp.*) | 7.85 | 20 |
| R112 | rough_lemon | Lemon (*Citrus jambhiri Lush.*) | 5.68 | 15 |
| R116 | rough_lemon | Lemon (*Citrus jambhiri Lush.*) | 7.86 | 20 |
| R66 | rough_lemon | Lemon (*Citrus jambhiri Lush.*) | 9.21 | 24 |
| R69 | macrophylla | Papeda (*C. ichangensis*) | 15.18 | 39 |
| R6 | volckameriana | Lemon (*C.* × *limon*) | 8.98 | 23 |
| R76 | macrophylla | Papeda (*C. ichangensis*) | 12.87 | 33 |
| R7 | volckameriana | Lemon (*C.* × *limon*) | 6.75 | 17 |
| R83 | volckameriana | Lemon (*C.* × *limon*) | 14.02 | 36 |
| R92 | volckameriana | Lemon (*C.* × *limon*) | 8.60 | 22 |
| R107 | carrizo | Citrange (*X Citroncirus sp.*) | 14.49 | 37 |
| R111 | carrizo | Citrange (*X Citroncirus sp.*) | 9.20 | 24 |
| R115 | carrizo | Citrange (*X Citroncirus sp.*) | 8.76 | 22 |
| R129 | carrizo | Citrange (*X Citroncirus sp.*) | 4.64 | 12 |
| R12 | carrizo | Citrange (*X Citroncirus sp.*) | 10.90 | 28 |
| R21 | carrizo | Citrange (*X Citroncirus sp.*) | 13.09 | 34 |
| R36 | C-35 | Citrange (*X Citroncirus sp.*) | 15.26 | 39 |
| R38 | carrizo | Citrange (*X Citroncirus sp.*) | 11.14 | 29 |
| R40 | carrizo | Citrange (*X Citroncirus sp.*) | 16.10 | 41 |
| R41 | carrizo | Citrange (*X Citroncirus sp.*) | 12.34 | 32 |
| R42 | carrizo | Citrange (*X Citroncirus sp.*) | 7.77 | 20 |
| R79 | carrizo | Citrange (*X Citroncirus sp.*) | 13.23 | 34 |
| R91 | carrizo | Citrange (*X Citroncirus sp.*) | 13.32 | 34 |
| R98 | C-35 | Citrange (*X Citroncirus sp.*) | 13.32 | 34 |
| R99 | C-35 | Citrange (*X Citroncirus sp.*) | 16.80 | 43 |
| R100 | carrizo | Citrange (*X Citroncirus sp.*) | 17.11 | 44 |

| | | | | |
|---|---|---|---|---|
| R113 | troyer | Citrange (*X Citroncirus sp.*) | 10.97 | 28 |
| R127 | carrizo | Citrange (*X Citroncirus sp.*) | 17.28 | 44 |
| R20 | carrizo | Citrange (*X Citroncirus sp.*) | 13.16 | 34 |
| R44 | carrizo | Citrange (*X Citroncirus sp.*) | 14.31 | 37 |
| R45 | carrizo | Citrange (*X Citroncirus sp.*) | 7.09 | 18 |
| R46 | carrizo | Citrange (*X Citroncirus sp.*) | 9.91 | 25 |
| R60 | carrizo | Citrange (*X Citroncirus sp.*) | 12.47 | 32 |
| R89 | carrizo | Citrange (*X Citroncirus sp.*) | 14.87 | 38 |
| R8 | carrizo | Citrange (*X Citroncirus sp.*) | 20.44 | 52 |
| R93 | carrizo | Citrange (*X Citroncirus sp.*) | 6.44 | 17 |
| R101 | carrizo | Citrange (*X Citroncirus sp.*) | 8.25 | 21 |
| R123 | carrizo | Citrange (*X Citroncirus sp.*) | 15.69 | 40 |
| R53 | cleopatra | Mandarin (*C. reticulata*) | 20.52 | 53 |
| R56 | carrizo | Citrange (*X Citroncirus sp.*) | 16.99 | 44 |
| R67 | volckameriana | Lemon (*C.* × *limon*) | 14.75 | 38 |
| R78 | volckameriana | Lemon (*C.* × *limon*) | 9.28 | 24 |
| R84 | citrus_tangerina | Mandarin (*C. reticulata*) | 11.61 | 30 |
| R94 | macrophylla | Papeda (*C. ichangensis*) | 8.71 | 22 |
| R108 | carrizo | Citrange (*X Citroncirus sp.*) | 7.83 | 20 |
| R39 | carrizo | Citrange (*X Citroncirus sp.*) | 14.68 | 38 |
| R51 | C-35 | Citrange (*X Citroncirus sp.*) | 10.91 | 28 |
| R61 | carrizo | Citrange (*X Citroncirus sp.*) | 7.61 | 20 |
